# Structure of human PIEZO1 and its slow inactivating channelopathy mutants

**DOI:** 10.1101/2024.07.14.603468

**Authors:** Yuanyue Shan, Xinyi Guo, Mengmeng Zhang, Meiyu Chen, Ying Li, Mingfeng Zhang, Duanqing Pei

## Abstract

PIEZO channels transmit mechanical force signals to cells, allowing them to make critical decisions during development and in pathophysiological conditions. Their fast/slow inactivation modes have been implicated in mechanopathologies, but remain poorly understood. Here, we report several near-atomic resolution cryo-EM structures of fast-inactivating wild-type human PIEZO1 (hPIEZO1) and its slow-inactivating channelopathy mutants with or without its auxiliary subunit MDFIC. Our results suggest that hPIEZO1 has a more flattened and extended architecture than curved mouse PIEZO1 (mPIEZO1). The multi-lipidated MDFIC subunits insert laterally into the hPIEZO1 pore module like mPIEZO1, resulting in a more curved and extended state. Interestingly, the high-resolution structures suggest that the pore lipids, which directly seal the central hydrophobic pore, may be involved in the rapid inactivation of hPIEZO1. While the severe hereditary erythrocytosis mutant R2456H significantly slows down the inactivation of hPIEZO1, the hPIEZO1-R2456H-MDFIC complex shows a more curved and contracted structure with an inner helix twist due to the broken link between the pore lipid and R2456H. These results suggest that the pore lipids may be involved in the mechanopathological rapid inactivation mechanism of PIEZO channels.

## Introduction

Cells rely on mechanosensitive (MS) channels, which rapidly convert force into ion flow, to sense environmental changes and make appropriate decisions throughout the prokaryotic and eukaryotic kingdoms. In bacteria, two major MS channels, the small-conductance mechanosensitive channel (MscS) ^1^ and large-conductance mechanosensitive channel (MscL) ^2^ channels, are responsible for bacterial force sensing. In contrast, in mammals, only the PIEZO ^3^, two-pore domain K^+^ channel (K2P) ^4^ and TMEM63 ^5,6^ channels have been identified as bona fide MS channels. Structural studies indicate that not all MS channels are constructively conserved, ranging from monomer ^7^ to heptamer ^8^, but they are likely to obey two putative principles of force-from-lipid (FFL) and force-from-filament (FFF) ^9,10^.Within the heptameric MscS channel in bacteria, three types of lipids, the pore lipids, the gatekeeper lipids, and the pocket lipids, sense and transduce force at corresponding positions ^11^. Similarly, the pseudo-tetrameric MS K2P channels also utilize at least three types of lipids to regulate channel activation ^12,13^. Particularly, pore lipids seal the channel pores and could be removed by the mechanical force upon activation of the MscS, MS K2P and TMEM63 channels ^7,11,12^. In the trimeric PIEZO channels, membrane curvature is likely associated with channel activation ^14–16^. Although the PIEZO channel also obeys the FFL principle^17^, the role of the lipids in PIEZO channel gating remains elusive.

PIEZOs from different species or in disparate native cells exhibit diverse inactivation kinetics. hPIEZO1 exhibits faster inactivation gating kinetics than mPIEZO1 under the same voltage condition^3,17,18^, so mPIEZO1 presents a native “GOF” state relative to hPIEZO1 regarding inactivation gating kinetics. Also, it has been reported recently that the MyoD-family inhibitor, MDFIC/MDFI, are auxiliary subunits of PIEZO channels to prolong inactivation and reduce mechanosensitivity^19^. Besides, some Hereditary erythrocytosis (HX) mutations exhibit slower inactivation gating kinetics than wildtype hPIEZO1. The slower inactivation may result in the higher open probability and delayed inactivation of hPIEZO1; thus, these mutations are considered gain of function (GOF) ^20^. HX, also known as inherited dehydrated stomatocytosis, is an autosomal dominant disorder that causes dehydration of red blood cells (RBCs), resulting in haemolytic anaemia. In addition to changes in gating kinetics, some of the HX mutations show alterations in response to osmotic pressure and in membrane protein trafficking ^21^. Interestingly, the populations with a mild GOF PIEZO1 allele are likely to be resistant to malaria infection, while some present a strong association with increased plasma iron ^22,23^. Although the inactivation phenotype of PIEZO channels is very susceptible to intrinsic and extrinsic factors, the molecular basis of their inactivation remains unclear.

Here, we report the structural basis of hPIEZO1 gating and inactivation based on the architecture of hPIEZO1 and its slow inactivating channelopathy mutants with or without its auxiliary subunit MDFIC at near-atomic resolution by cryo-EM. Our high-resolution structures support a model in which the pore lipids directly seal the central hydrophobic pore and involve fast inactivation of hPIEZO1. This model also provides a mechanistic understanding of the severe hereditary erythrocytosis mutant R2456H with a more curved and contracted structure with an inner helix twist due to the broken link between the pore lipid and R2456H.

## Results

### Overall structure of the full-length hPIEZO1

hPIEZO1 is known to have faster inactivation gating kinetics than mPIEZO1 under the same voltage condition^3,18,20^. The structural basis of this difference remains unresolved. To resolve this, we synthesized a codon-optimized full-length hPIEZO1 with a C-terminal GFP and flag tag into the pBM vector ^24^ and this construct exhibits similar mechanosensitivity and fast inactivation gating properties as previously reported by whole-cell poking assay (Fig. S2a). We overexpressed hPIEZO1 in 12 liters of HEK293F suspension cells, extracted the protein with lauryl maltose neopentyl glycol (LMNG) and purified it in a digitonin environment. The symmetric peak of the full-length hPIEZO1 suggests the protein is homogenous. The smeared bands around 300 kDa on SDS-PAGE indicate the potential presence of post-translational modifications (PTMs) and the lower bands may be the binding partners or degraded fragments of hPIEZO1 (Fig. S1a-b).

The purified hPIEZO1 protein was subjected to the standard cryo-EM workflow, resulting in a cryo-EM density map at a total resolution of 3.3 Å from approximately 87K particles (Fig. S3a-e) that allowed us to build a near-atomic model of hPIEZO1 (Fig. S4). Among the nine transmembrane helix units (THUs) containing four transmembrane helices for each predicted by primary sequences, the two N-terminal ones are not visible in the cryo-EM density map. The rest of the THUs, along with the pore module consisting of an anchor, an outer helix (OH), an inner helix (IH), and a cap domain, are resolved (Fig.1a). Compared to the anchor, OH and IH, the resolution of the cap domain is slightly lower, which may be due to the flexibility of the cap domain. On the intracellular side, a long beam helix supports the THU7-9 and pore modules. Each hPIEZO1 subunit has a curved blade structure and the three subunits form the trimeric PIEZO channel, exhibiting a bowl-like shape. The hPIEZO1 density map, filtered with a low pass filter, shows a disk approximately 30 nm wide with a horn-like density in the peripheral blade region, likely due to the PTMs such as glycosylation. Interestingly, we see a large density below the central pore module at the intracellular side, which may be potential auxiliary subunits, consistent with the lower molecular weight bands on SDS-PAGE (Fig. 1b-g).

**Fig. 1.**
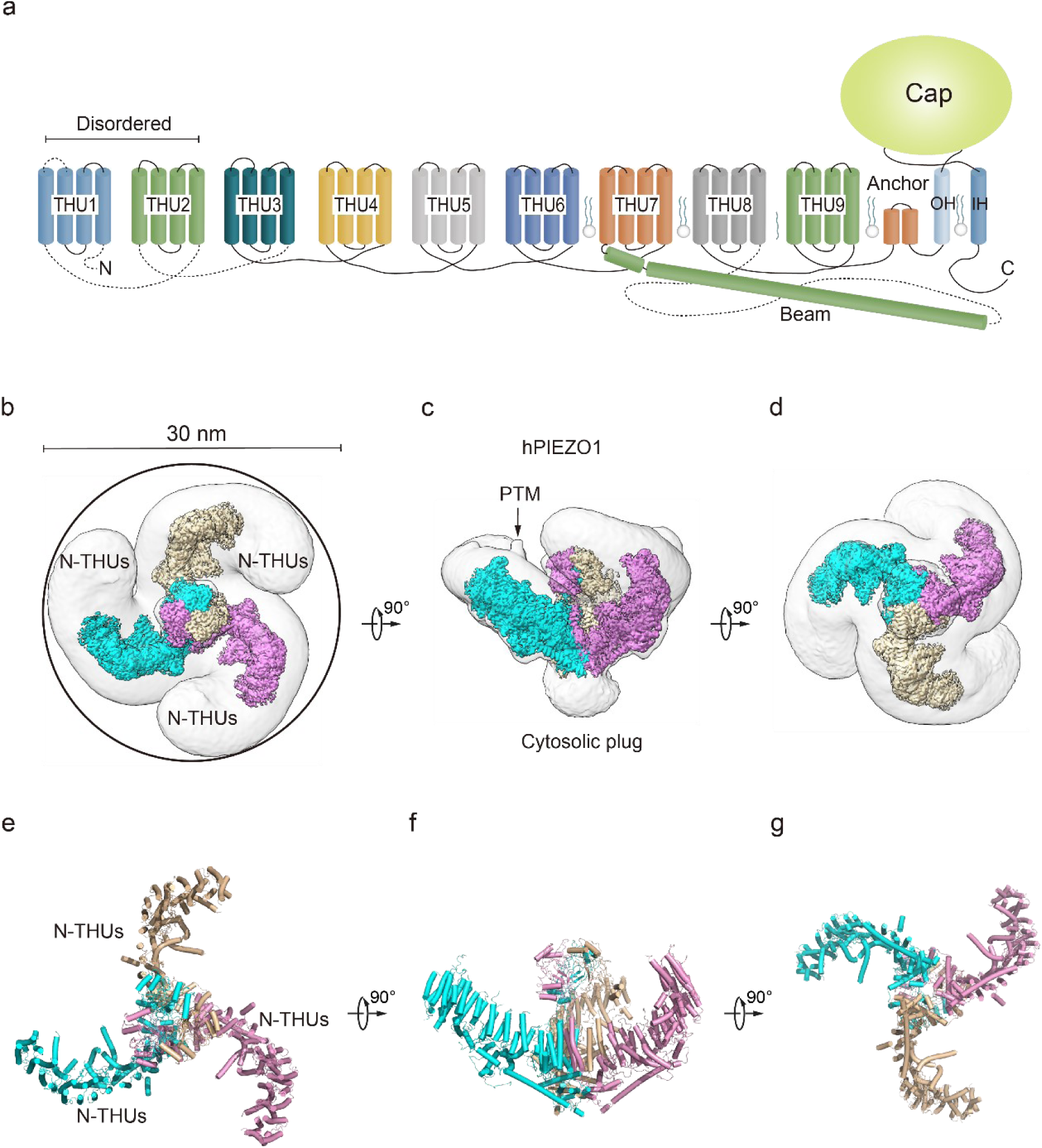
Structure of full-length human PIEZO1 channel **a,** 38-TM topology model of a single hPIEZO1 subunit. The 9 THUs, a long beam helix, an anchor domain, and two pore module helices (OH and IH, respectively). Each THU contains four transmembrane helices. THU1 and THU2 are likely disordered and therefore not visible in the hPIEZO1 cryo-EM density map. **b**, The 3.3 Å cryo-EM density map of hPIEZO1, viewed from the top. The density of each single subunit is colored in cyan, wheat and pink, respectively. Around 30 nm digitonin disk is shown as grey density by low pass filtered. **c**, The 3.3 Å cryo-EM density map of hPIEZO1, viewed from the side. The density of each single subunit is colored in cyan, wheat and pink, respectively. The potential PTM at the THUs region is indicated. A flexible density binds to the cytosolic plug, which may stand for an additional hPIEZO1 auxiliary. **d**, The 3.3 Å cryo-EM density map of hPIEZO1, viewed from the bottom. The density of each single subunit is colored in cyan, wheat and pink, respectively. **e**, The cartoon model of hPIEZO1, viewed from the top. Each single subunit is colored in cyan, wheat and pink, respectively. **f**, The cartoon model of hPIEZO1, viewed from the side. Each single subunit is colored in cyan, wheat and pink, respectively. **g**, The cartoon model of hPIEZO1, viewed from the bottom. Each single subunit is colored in cyan, wheat and pink, respectively.

### hPIEZO1 is more flattened and extended than mPIEZO1

Several studies using high-speed atomic force microscopy (HS-AFM) ^14^, cryo-EM^16^ and 3D interferometric photoactivation localization microscopy (iPALM)^15^ suggest that PIEZO blades may receive mechanical force through sensing membrane curvature. The reshaping of the curvature of the PIEZO conducts force to the pore region, which may result in the channel pore opening and ion flow. Structural comparison of hPIEZO1 and curved mPIEZO1 shows that hPIEZO1 presents a more flattened structure from the side view (Fig. 2a). The blades of hPIEZO1 are about 5 Å down towards the cytoplasmic side than those of mPIEZO1. Meanwhile, the cap domain of hPIEZO1 is slightly upwards to the extracellular side (Fig. 2a). In the top view, the distal blades rotate counterclockwise about 22 Å compared to mPIEZO1 (Fig. 2b). Therefore, hPIEZO1 appears more extended than the curved mPIEZO1. Meanwhile, hPIEZO1 still maintains a sizeable curved state compared to the flattened mPIEZO1 structure. From the top view, the two structures are very similar (Fig. 2c-d). The pore radius of hPIEZO1 is between that of the curved and flattened mPIEZO1. Therefore, the curved hPIEZO1 may still represent a non-conducting state (Fig. 2e-h), and the differences in curvature and pore radius may be implicated in channel gating properties of hPIEZO1 and mPIEZO1.

**Fig. 2.**
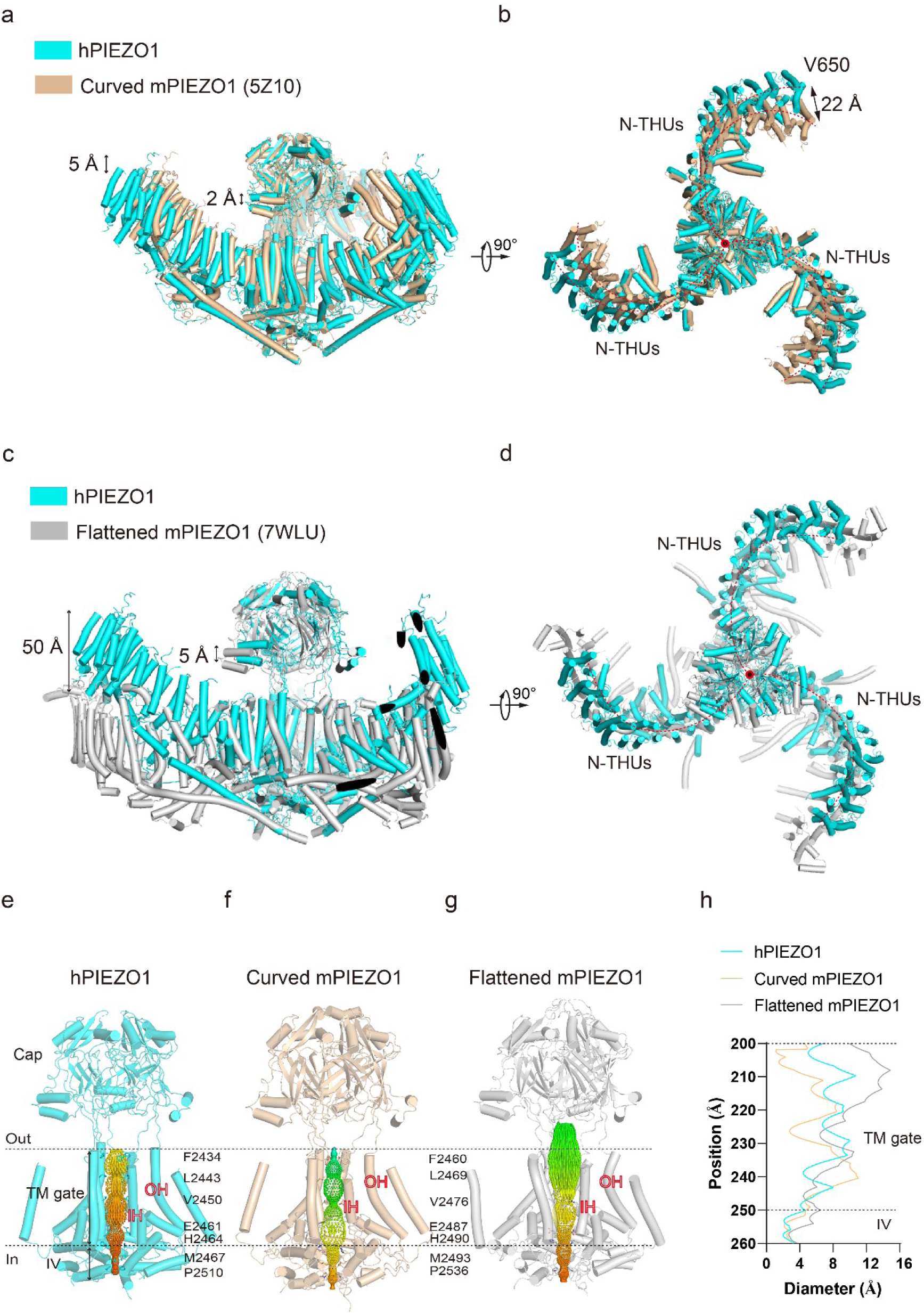
Structural comparison of hPIEZO1 and mPIEZO1 **a,** Structural comparison of hPIEZO1 (this study) and curved mPIEZO1 (5Z10), viewed from the side. The distance of the distal blade between curved mPIEZO1 and hPIEZO1 is measured as 5 Å, and the distance of the cap between curved mPIEZO1 and hPIEZO1 is measured as 2 Å. **b**, Structural comparison of hPIEZO1(this study) and curved mPIEZO1 (5Z10), viewed from the top. The distance between the V650 residue in hPIEZO1 and the corresponding residue in curved mPIEZO1 is around 22 Å; therefore, the blades of hPIEZO1 present a contracted state from the top view. The blade center lines are shown as red dashed lines. **c**, Structural comparison of hPIEZO1 (this study) and flattened mPIEZO1 (7WLU), viewed from the side. Compared to the flattened mPIEZO1, the mimic membrane curvature of hPIEZO1 is between the curved and flattened mPIEZO1. **d**, Structural comparison of hPIEZO1(this study) and curved mPIEZO1 (5Z10), viewed from the top. hPIEZO1 presents a similar extended state to the flattened mPIEZO1. **e**, The cartoon model of the hPIEZO1 pore module with calculated pore. The restricted residues are labeled. **f**, The cartoon model of curved mPIEZO1 pore module with calculated pore. The restricted residues are labeled. **g**, The cartoon model of flattened mPIEZO1 pore module with calculated pore. **h**, The calculated pore diameter of hPIEZO1 (cyan), curved mPIEZO1 (wheat) and flattened mPIEZO1(grey) along the z axis.

### GOF channelopathy mutants, hPIEZO1-A1988V, hPIEZO1-E756del and hPIEZO1-R2456H, are structurally unstable

The success in solving the hPIEZO1 structure encouraged us to probe the structural basis of channelopathy. We selected the mild GOF mutants hPIEZO1-A1988V and hPIEZO1-E756del, familiar in African populations with a slightly longer inactivation time, and the more severe R2456H mutant with a much longer inactivation time ^20,23^. However, when we tried to solve their structures based on the same method, i.e., overexpressing and purifying these three channelopathy mutants in HEK293F suspension cells, extracting them with LMNG detergent and purifying them in a digitonin environment (Fig. S1c-e), we can only obtain three channelopathy mutant proteins and their raw cryo-EM images. Still, we failed to achieve satisfactory 2D class averages for them (Fig. S5-6 and S7a-b). As a result, we could not generate high-resolution 3D density maps of hPIEZO1-E756del and hPIEZO1-R2456H. We then tried and obtained almost twice as many hPIEZO1-A1988V images as the wild type but only achieved a 3.7 Å resolution cryo-EM density map (Fig. S7). Yet, the resulting overall structure is similar to the wild type (Fig. S8). Based on these results, these three hPIEZO1 channelopathy GOF mutants may be more structurally unstable than the wild type.

### The structure of the hPIEZO1-MDFIC complex

MDFIC, a MyoD family inhibitor protein, has recently been identified as an auxiliary subunit of piezo channels capable of decelerating channel inactivation and attenuating channel mechanosensitivity ^19^. To solve the structural basis of MDFIC on hPIEZO1, we co-expressed it with hMDFIC in HEK293F and purified the complex (Fig. S1a, S1f). We obtained a 3.0 Å resolution cryo-EM density map of the hPIEZO1-MDFIC complex (Fig. S9). The overall cryo-EM density map is similar to hPIEZO1 alone with a disc of about 30 nm (Fig. 3a-f), but allowed us to validate the MDFIC density with multi-lipidated cysteines on the C-terminal amphipathic helix (Fig. S12). This C-terminal helix inserted laterally into the pore module where the exact position of mPIEZO1 is found clearly (Fig.3a-f). Interestingly, while MDFIC does not significantly alter the curvature of mPIEZO1, it induces a more curved and contracted architecture in hPIEZO1 from the side and top views, respectively (Fig.3g-h). Combining with the similar effect of MDFIC on mPIEZO1^25^ and hPIEZO1 (Fig. S2), we suspect that MDFIC decelerates hPIEZO1 inactivation and weakens its mechanosensitivity by remodeling the pore module and exerting more blade curvature.

**Fig. 3.**
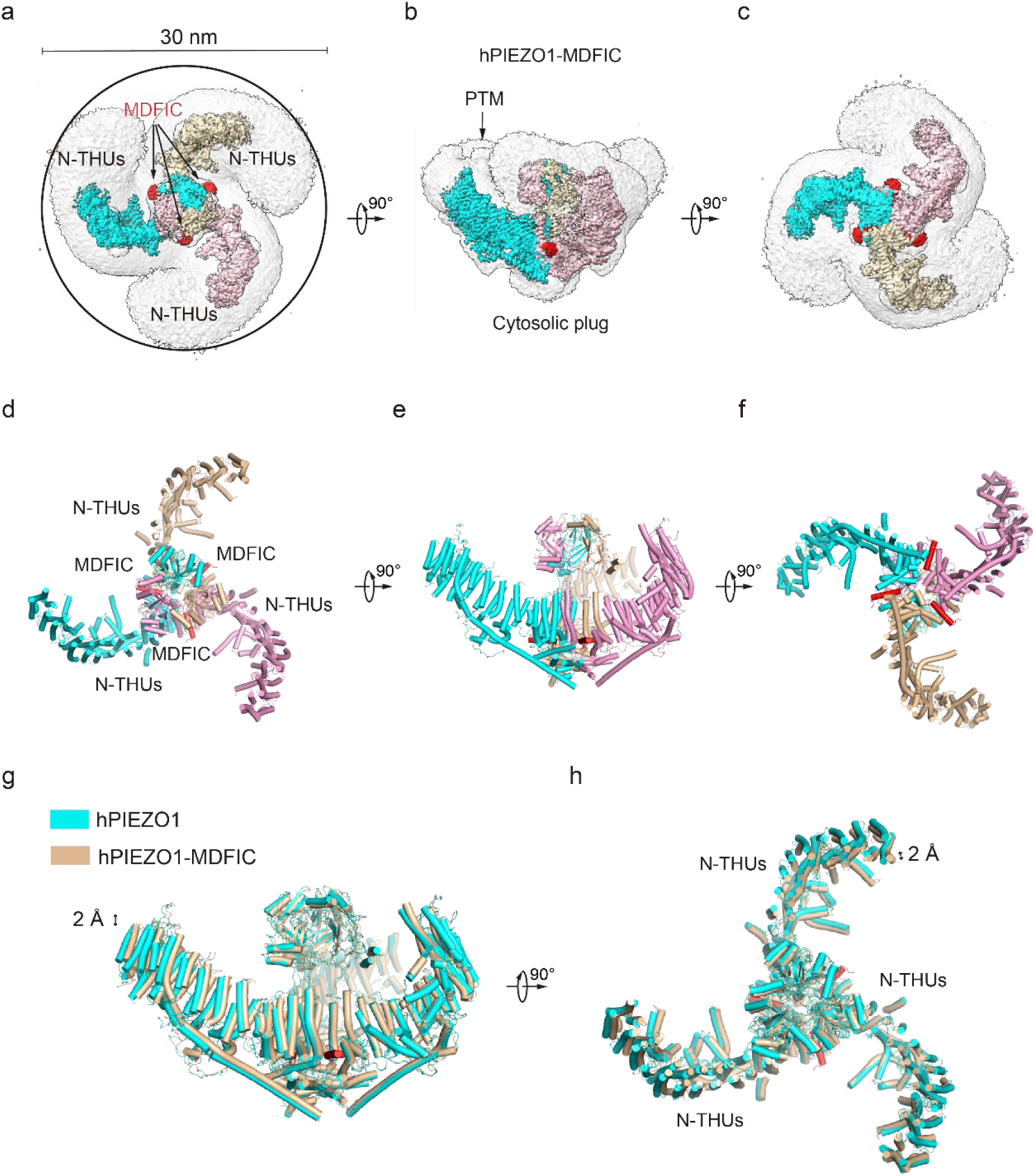
Structure of full-length human PIEZO1-MDFIC complex **a**, The 3.0 Å cryo-EM density map of hPIEZO1-MDFIC, viewed from the top. The density of each single subunit is colored in cyan, wheat and pink, respectively. The density of the three C-terminal helices of MDFIC is colored in red. Around 30 nm digitonin disk is shown as grey density by low pass filtered. **b**, The 3.0 Å cryo-EM density map of hPIEZO1-MDFIC, viewed from the side. The density of each single subunit is colored in cyan, wheat and pink, respectively. The density of the three C-terminal helices of MDFIC is colored in red. The potential PTM at the THUs region is indicated. A flexible density binds to the cytosolic plug, which may stand for an additional hPIEZO1 auxiliary subunit. **c**, The 3.0 Å cryo-EM density map of hPIEZO1-MDFIC, viewed from the bottom. The density of each single subunit is colored in cyan, wheat and pink, respectively. The density of three C-terminal helices of MDFIC is colored in red. **d**, The cartoon model of hPIEZO1-MDFIC, viewed from the top. Each single subunit is colored in cyan, wheat and pink, respectively. Three C-terminal helices of MDFIC are colored in red. **e**, The cartoon model of hPIEZO1-MDFIC, viewed from the side. Each single subunit is colored cyan, wheat and pink, respectively. Three C-terminal helices of MDFIC are colored in red. **f**, The cartoon model of hPIEZO1-MDFIC, viewed from the bottom. Each single subunit is colored in cyan, wheat and pink, respectively. Three C-terminal helices of MDFIC are colored in red. **g,** Structural comparison of hPIEZO1 and hPIEZO1-MDFIC, viewed from the side. The motion of the distal blade between hPIEZO1 and hPIEZO1-MDFIC is around 2 Å from the side view. **h**, Structural comparison of hPIEZO1 and hPIEZO1-MDFIC, viewed from top. The motion of the distal blade between hPIEZO1 and hPIEZO1-MDFIC is around 2 Å from the top view.

### MDFIC stabilizes GOF channelopathy hPIEZO mutant complexes

The gating kinetics change of MDFIC on hPIEZO1 further encouraged us to test if it can stabilize the unstable channelopathy mutants hPIEZO1-A1988V, hPIEZO1-E756del and hPIEZO1-R2456H (Fig. 4a). Indeed, electrophysiological studies showed that co-expression of these channelopathy mutants with MDFIC resulted in significantly reduced mechanosensitivity and inactivation rate (Fig. S2). We then co-expressed these hPIEZO1 mutants with MDFIC in HEK293F suspension cells and purified the resulting complex using the same method (Fig. S1g-i). Surprisingly, with standard cryo-EM analysis, we obtained high-quality 2D class averages for hPIEZO1-A1988V-MDFIC and hPIEZO1-E756del-MDFIC. Cryo-EM density maps were obtained for hPIEZO1-A1988V-MDFIC and hPIEZO1-E756del-MDFIC with overall resolutions of 3.1 and 2.8 Å, respectively (Fig. S10- 11). The densities of the multi-lipidated C-terminal amphipathic helix of MDFIC are present in the hPIEZO1-A1988V-MDFIC and hPIEZO1-E756del-MDFIC maps. For hPIEZO1-R2456H-MDFIC, the 2D class averages show a clear trimer state (Fig. S13b). The resolution of hPIEZO1-R2456H- MDFIC was improved to 4.6 Å with 16K particles, allowing to build a relatively accurate model of the transmembrane helices based on the hPIEZO1-MDFIC model (Fig. S13-14). These results further suggest that MDFIC stabilizes the GOF channelopathy mutants structurally.

**Fig. 4.**
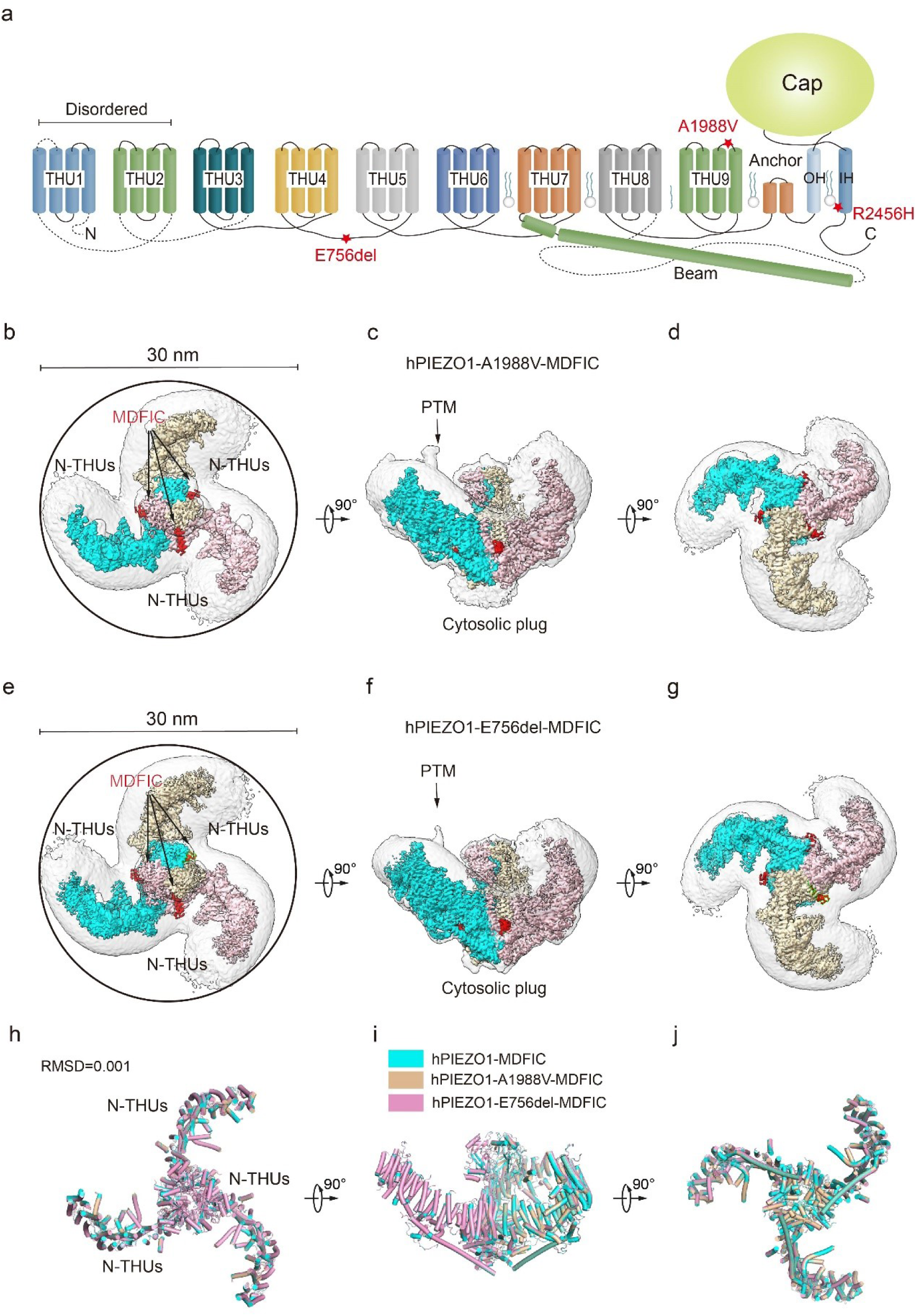
Structure of human PIEZO1-A1988V-MDFIC and hPIEZO1-E756del-MDFIC complex a, 38-TM topology model of a single hPIEZO1 subunit. The E756del mutation is located at the THU4 and THU5 linker, the A1988V mutation is located at the THU9 linker and the R2456H mutation is located at the IH of the pore module. **b**, The 3.1 Å cryo-EM density map of hPIEZO1-A1988V-MDFIC, viewed from the top. The density of each single subunit is colored in cyan, wheat and pink, respectively. The density of the three C-terminal helices of MDFIC is colored in red. Around 30 nm digitonin disk is shown as grey density by low pass filtered. **c**, The 3.1 Å cryo-EM density map of hPIEZO1-A1988V-MDFIC, viewed from the side. The density of each single subunit is colored in cyan, wheat and pink, respectively. The density of the three C-terminal helices of MDFIC is colored in red. The potential PTM at the THUs region is indicated. A flexible density binds to the cytosolic plug, which may represent an additional hPIEZO1 auxiliary. **d**, The 3.1 Å cryo-EM density map of hPIEZO1-A1988V-MDFIC, viewed from the bottom. The density of each single subunit is colored in cyan, wheat and pink, respectively. **e**, The 2.8 Å cryo-EM density map of hPIEZO1-E756del-MDFIC, viewed from the top. The density of each single subunit is colored in cyan, wheat and pink, respectively. The density of the three C-terminal helices of MDFIC is colored in red. Around 30 nm digitonin disk is shown as grey density by low pass filtered. **f**, The 2.8 Å cryo-EM density map of hPIEZO1-E756del-MDFIC, viewed from the side. The density of each single subunit is colored in cyan, wheat and pink, respectively. The density of the three C-terminal helices of MDFIC is colored in red. The potential PTM at the THUs region is indicated. A flexible density binds to the cytosolic plug, which may represent an additional hPIEZO1 auxiliary. **g**, The 2.8 Å cryo-EM density map of hPIEZO1-E756del-MDFIC, viewed from the bottom. The density of each single subunit is colored in cyan, wheat and pink, respectively. **h**, Superimposed cartoon models of hPIEZO1-MDFIC (cyan), hPIEZO1-A1988V-MDFIC (wheat) and hPIEZO1-E756del-MDFIC (pink), viewed from the top. The RSMD is around 0.001, indicating that they are almost identical. **i**, Superimposed cartoon models of hPIEZO1-MDFIC (cyan), hPIEZO1-A1988V-MDFIC (wheat) and hPIEZO1-E756del-MDFIC (pink), viewed from the side. **j**, Superimposed cartoon models of hPIEZO1-MDFIC (cyan), hPIEZO1-A1988V-MDFIC (wheat) and hPIEZO1-E756del-MDFIC (pink), viewed from the bottom.

### hPIEZO1-R2456H-MDFIC with more curved blades but more extended pore

The unexpected higher resolution maps of hPIEZO1-A1988V-MDFIC and hPIEZO1-E756del- MDFIC reveal a bowl-like disc of about 30 nm for both complexes (Fig. 4h-j). On the other hand, the hPIEZO1-R2456H-MDFIC shows a smaller disc of 25 nm compared to the 30 nm for the wild type and the other two mutants (Fig. 5a-f). The smaller disc is mainly due to the more significant contraction of the blade arms and a more curved state compared to the wild-type hPIEZO1-MDFIC in the side view (Fig. 5g-h). Surprisingly, the IH of hPIEZO1-R2456H-MDFIC showed a significant twist of ∼35 degrees compared to wild-type hPIEZO1-MDFIC (Fig. 6g-h). The coiled-coil shape of the IH pore results in a more dilated pore on the extracellular side (Fig. 5i-l), suggesting that unlike the mild GOF E756del and A1988V mutations present in the typical African population and located in the THUs region, the more severe mutation R2456H, which is located in the IH, not only leads to blade reshaping but also results in pore expansion (Fig. 6d-h).

**Fig. 5.**
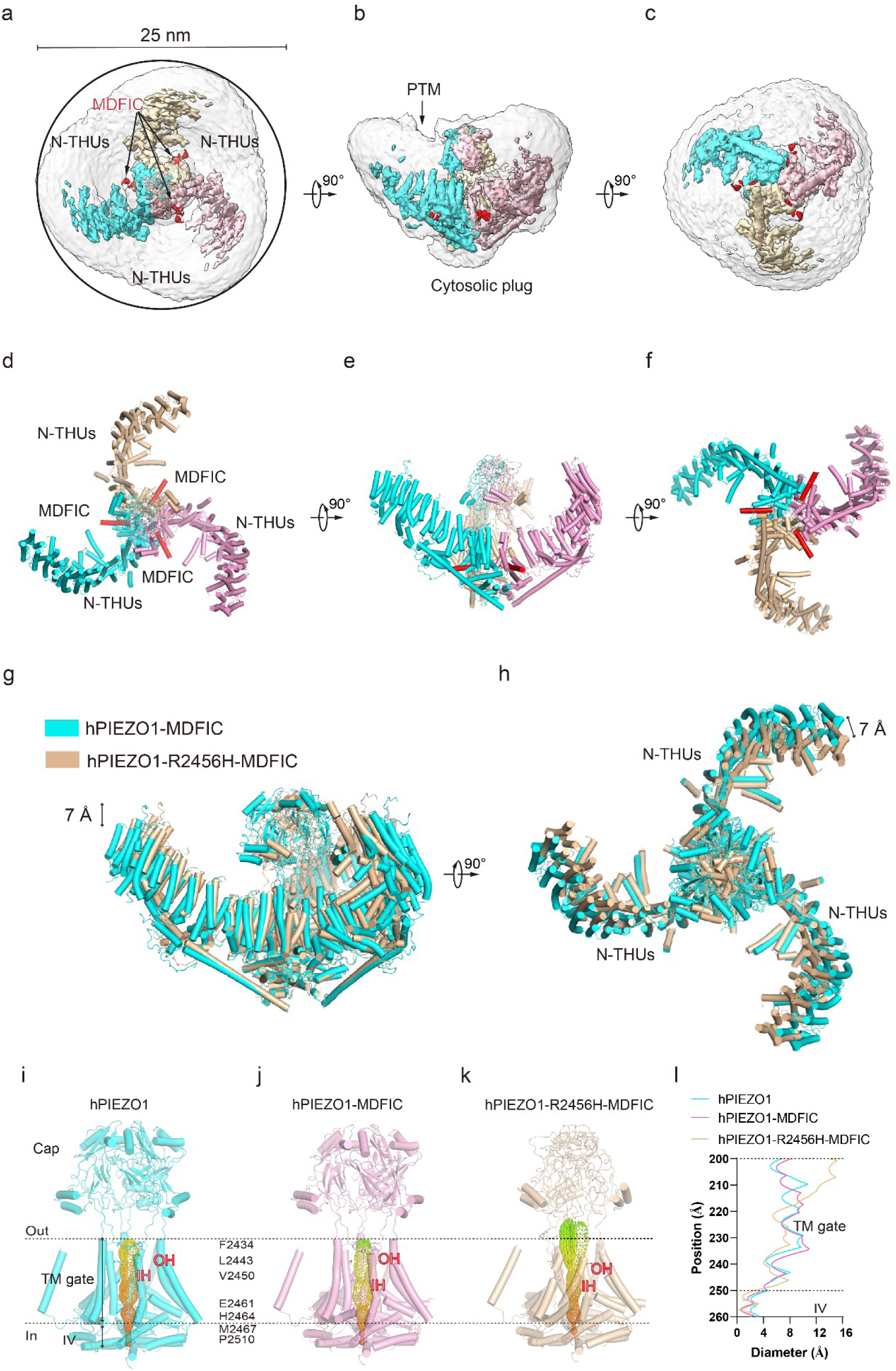
Structure of human PIEZO1-R2456H-MDFIC complex **a**, The 4.5 Å cryo-EM density map of hPIEZO1-R2456H-MDFIC, viewed from the top. The density of each single subunit is colored in cyan, wheat and pink, respectively. The density of the three C-terminal helices of MDFIC is colored in red. Around 25 nm digitonin disk of hPIEZO1-R2456H- MDFIC is shown as grey density by low pass filtered, smaller than the wildtype hPIEZO1-MDFIC. **b**, The 4.5 Å cryo-EM density map of hPIEZO1-R2456H-MDFIC, viewed from the side. The density of each single subunit is colored in cyan, wheat and pink, respectively. The density of the three C-terminal helices of MDFIC is colored in red. The potential PTM at the THUs region is indicated. A flexible density binds to the cytosolic plug, which may represent an additional hPIEZO1 auxiliary. **c**, The 4.5 Å cryo-EM density map of hPIEZO1-R2456H-MDFIC, viewed from the bottom. The density of each single subunit is colored in cyan, wheat and pink, respectively. The density of the three C-terminal helices of MDFIC is colored in red. **d**, The cartoon model of hPIEZO1-R2456H-MDFIC, viewed from the top. Each single subunit is colored in cyan, wheat and pink, respectively. Three C-terminal helices of MDFIC are colored in red. **e**, The cartoon model of hPIEZO1-R2456H-MDFIC, viewed from the side. Each single subunit is colored in cyan, wheat and pink, respectively. Three C-terminal helices of MDFIC are colored in red. **f**, The cartoon model of hPIEZO1-R2456H-MDFIC, viewed from the bottom. Each single subunit is colored in cyan, wheat and pink, respectively. Three C-terminal helices of MDFIC are colored in red. **g,** Structural comparison of hPIEZO1-R2456H-MDFIC and hPIEZO1-MDFIC, viewed from the side. The motion of the distal blade between hPIEZO1-R2456H-MDFIC and hPIEZO1-MDFIC is around 7 Å from the side view. **h**, Structural comparison of hPIEZO1-R2456H-MDFIC and hPIEZO1-MDFIC, viewed from the top. The motion of the distal blade between hPIEZO1-R2456H-MDFIC and hPIEZO1-MDFIC is around 7 Å from the top view. **i**, The cartoon model of hPIEZO1 pore module with calculated pore. The restricted residues are labeled. **j**, The cartoon model of hPIEZO1-MDFIC pore module with calculated pore. **k**, The cartoon model of hPIEZO1-R2456H-MDFIC pore module with calculated pore. **l**, The calculated pore diameter of hPIEZO1 (cyan), hPIEZO1-MDFIC (pink) and flattened hPIEZO1-R2456H-MDFIC (wheat) along the z-axis. The hPIEZO1-R2456H-MDFIC presents a more extended extracellular side pore.

**Fig. 6.**
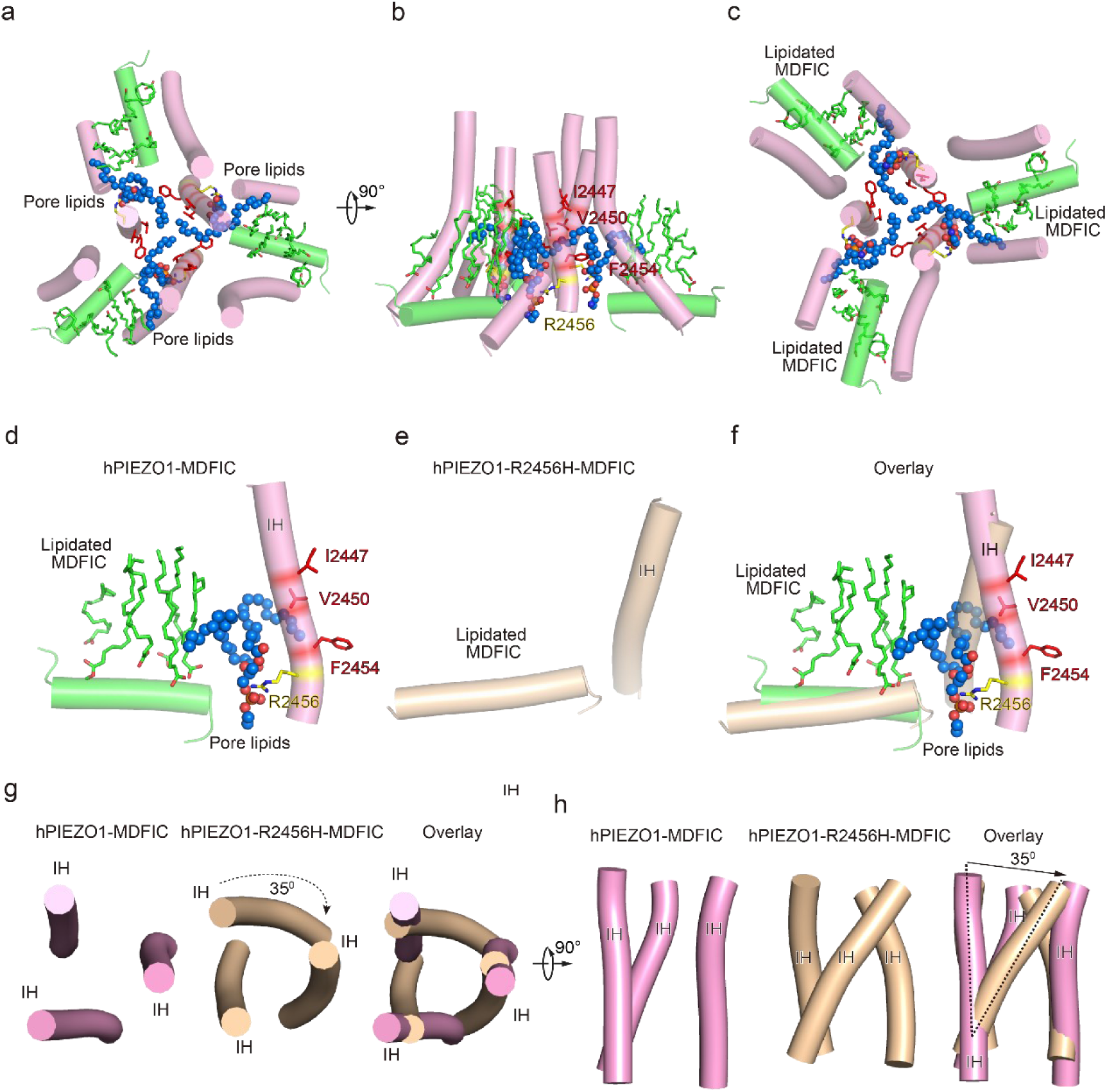
The pore module, multi-lipidated MDFIC and pore lipid interaction and the pore remodeling of the HX pathogenesis R2456H mutant **a**, The cartoon model of hPIEZO1-MDFIC/hPIEZO1-A1988V-MDFIC/hPIEZO1-E756del- MDFIC pore module, viewed from the top. The marine spheres present the pore lipids. **b**, The cartoon model of hPIEZO1-MDFIC/hPIEZO1-A1988V-MDFIC/hPIEZO1-E756del- MDFIC pore module, viewed from the side. The hydrophobic pore region formed by I2447, V2450 and F2454 is marked by red. The R2456 on the lateral side of the inner helix of the pore is marked by yellow. **c**, The cartoon model of hPIEZO1-MDFIC/hPIEZO1-A1988V-MDFIC/hPIEZO1-E756del-MDFIC pore module, viewed from the bottom. The multi-lipidated MDFIC subunits are colored in green and marked. One hydrophobic fatty acid chain of the pore lipid occupies the hydrophobic pore region. Another hydrophobic fatty acid chain of the pore lipid interacts with the fatty acid chains of the MDFIC-covalently-linked lipids. In contrast, the hydrophilic head of the pore lipid directly interacts with the R2456 on the lateral side of the inner helix of the pore, thus forming a stable hPIEZO1- multi-lipidated MDFIC-pore lipid complex. The pore lipids seal the pore and prevent ion flow. **d**, The cartoon model of hPIEZO1-MDFIC/hPIEZO1-A1988V-MDFIC/hPIEZO1-E756del- MDFIC IH, pore lipid and multi-lipidated MDFIC interactions. The multi-lipidated MDFIC subunits are colored in green. The marine spheres present the pore lipids. The hydrophobic pore region formed by I2447, V2450 and F2454 is marked by red. The R2456 on the lateral side of the inner helix of the pore is marked by yellow. **e**, The cartoon model of twisted IH and MDFIC in hPIEZO1-R2456H-MDFIC. **f**, The overlay cartoon model of IH and MDFIC in hPIEZO1-MDFIC/ hPIEZO1-A1988V-MDFIC/hPIEZO1-E756del-MDFIC and hPIEZO1-R2456H-MDFIC. **g**, The cartoon model of TM pore in hPIEZO1-MDFIC/ hPIEZO1-A1988V-MDFIC/ hPIEZO1- E756del-MDFIC and hPIEZO1-R2456H-MDFIC, viewed from the top. The TM pore of hPIEZO1- R2456H-MDFIC presents a twisted and extended state. **h**, The cartoon model of TM pore in hPIEZO1-MDFIC/ hPIEZO1-A1988V-MDFIC/ hPIEZO1- E756del-MDFIC and hPIEZO1-R2456H-MDFIC, viewed from the side. The TM pore of hPIEZO1- R2456H-MDFIC presents a twisted and extended state.

### R2456 anchors a lipid at the pore module

Then, we wished to build a pore module based on the solved structures. We were surprised to find an apparent lipid density in wild-type hPIEZO1-MDFIC, hPIEZO1-A1988V-MDFIC and even more evident in the 2.8 Å resolution map of hPIEZO1-E756del-MDFIC in the pore module, where one of its hydrophobic fatty acid tails inserts into the hydrophobic pore formed by I2447, V2450 and F2454, thus sealing the pore to prevent ion flow (Fig. 6a-c). Even more surprisingly, the hydrophilic phosphate head interacts directly with R2456, a residue with the more severe form of GOF mutation, on the side of the inner helix of the pore. In addition, another fatty acid chain of the pore lipid interacts with the acyl chains of the covalently linked MDFIC lipids, forming a stable hPIEZO1-multi-lipidated MDFIC-pore lipid complex (Fig. 6a-c). We also carefully checked the corresponding pore lipid configuration in the 3.3 Å resolution map of hPIEZO1 and the 3.8 Å resolution map of hPIEZO1-A1988V without the multi-lipidated MDFIC auxiliary subunit and found that in wild-type hPIEZO1, similar pore lipids also insert into the hydrophobic pore interacting with R2456 (Fig. 7a and 7f). However, in the mild GOF hPIEZO1-A1988V, which has a slower inactivation rate compared to wild-type hPIEZO1, the acyl chains of the pore lipid are retracted from the central hydrophobic pore to the side of the IH, although the hydrophilic phosphate group heads still interact with R2456 (Fig. 7b and 7f). These results reveal a critical role of R2456 in anchoring lipids at the pore module.

**Fig. 7.**
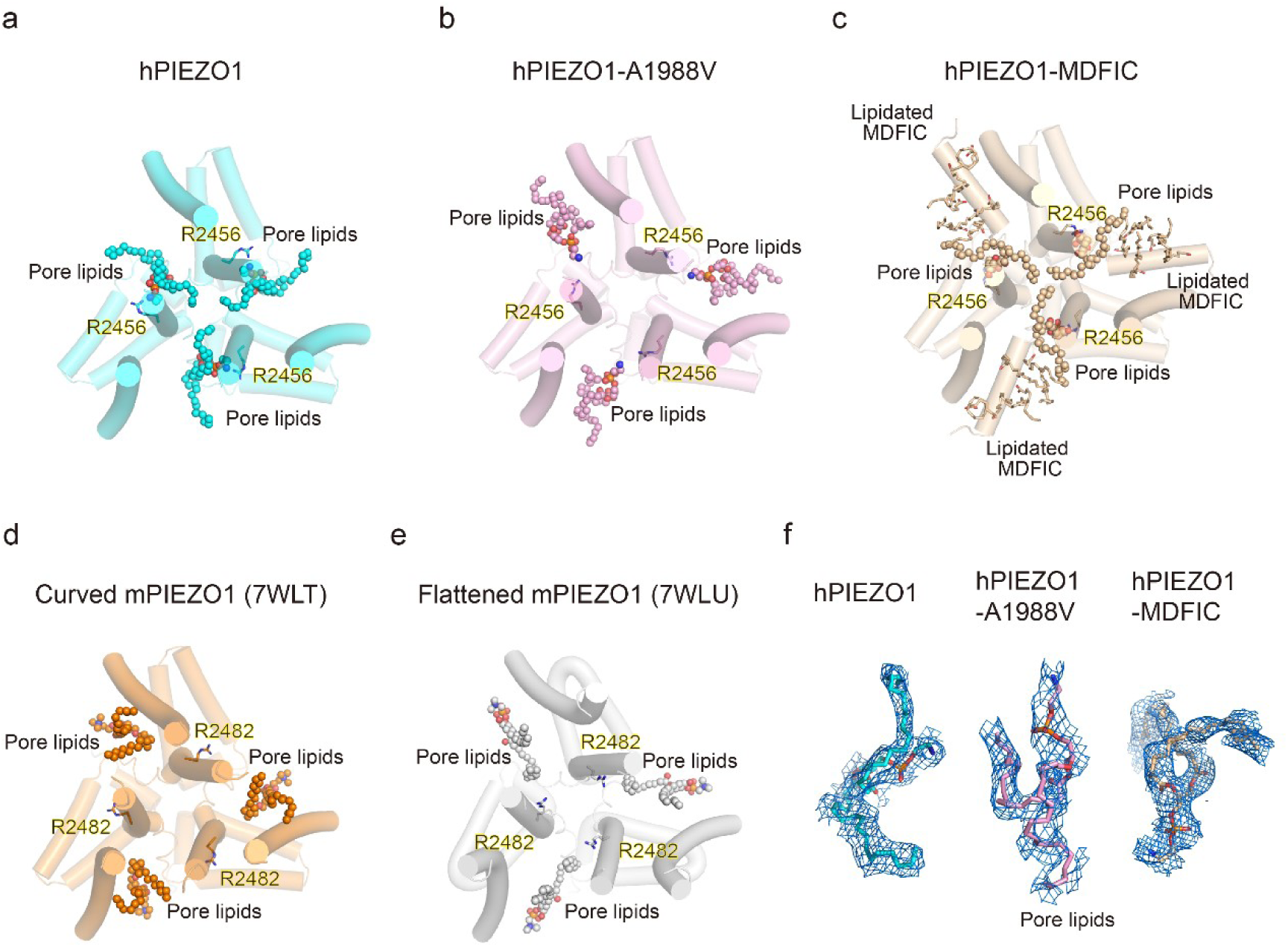
Occupancy pattern of pore lipids in hPIEZO1 underlying fast inactivation **a**, Top view of hPIEZO1 pore domain. The pore helices are shown as cylinders, and the pore lipid sealing the hydrophobic pore region is shown as spheres. The residue R2456s in inner helices, interacting with pore lipids, are labeled. **b**, Top view of hPIEZO1-A1988V pore domain. Pore lipids are retracted from the hydrophobic pore region. **c,** Top view of hPIEZO1-MDFIC pore domain. Lipidated-MDFICs are shown as sticks. **d**, Top view of curved mPIEZO1 (PDB: 7WLT). The residues R2482s, responding to R2456 in hPIEZO1, are labeled. Pore lipids are modeled in the latency side of the IH pore. **e**, Top view of flattened mPIEZO1 (PDB: 7WLU). Pore lipids are also located in the latency side of the IH pore. **f**, Cryo-EM density of pore lipids in WT hPIEZO1 (cyan), hPIEZO1-A1988V (pink) and hPIEZO1- MDFIC (brown). The cryo-EM density is shown as blue mesh.

### Pore lipids may involve the fast inactivation of hPIEZO1

Our results support a putative model that the hydrophobic acyl chain tails of the pore lipids insert into the hydrophobic pore region and seal the pore (Fig. 8a). Consistently, the slower inactivating hPIEZO1-A1988V mutant has the same hydrophobic acyl chain tails retracted from the hydrophobic pore region, implying that the pore lipids involve the fast inactivation of hPIEZO1 (Fig. 8a and 8c). The evidence supporting this model is based on previous electrophysiological functional studies that substitution of the hydrophobic pore, formed by I2447, V2450, and F2454, with a hydrophilic pore prolongs the inactivation time for both PIEZO1 and PIEZO2 channels ^17^. More robust evidence comes from the HX channelopathy mutant R2456H, wherein the interaction between the hydrophilic phosphate group head and R2456 is disrupted, which leads to remodeling of the blade and pore module (Fig. 6d-h), thus significantly prolonging the inactivation time. These results suggest that the pore lipids involve the fast inactivation of hPIEZO1 (Fig. 8d). Consistently, in curved and flattened mPIEZO1 structures ^16^, pore lipids are occupying the similar lateral side of the pore, but not sealing the pore-like the hPIEZO1-A1988V mutant (Fig. 7d-e and 8c). Similarly, we suspect that the curved and flattened mPIEZO1 structures are not sealed by pore lipid and thus may not adopt the deep inactivation state, consistent with electrophysiological results showing a slower inactivation manner of mPIEZO1^3,18^.

**Fig. 8.**
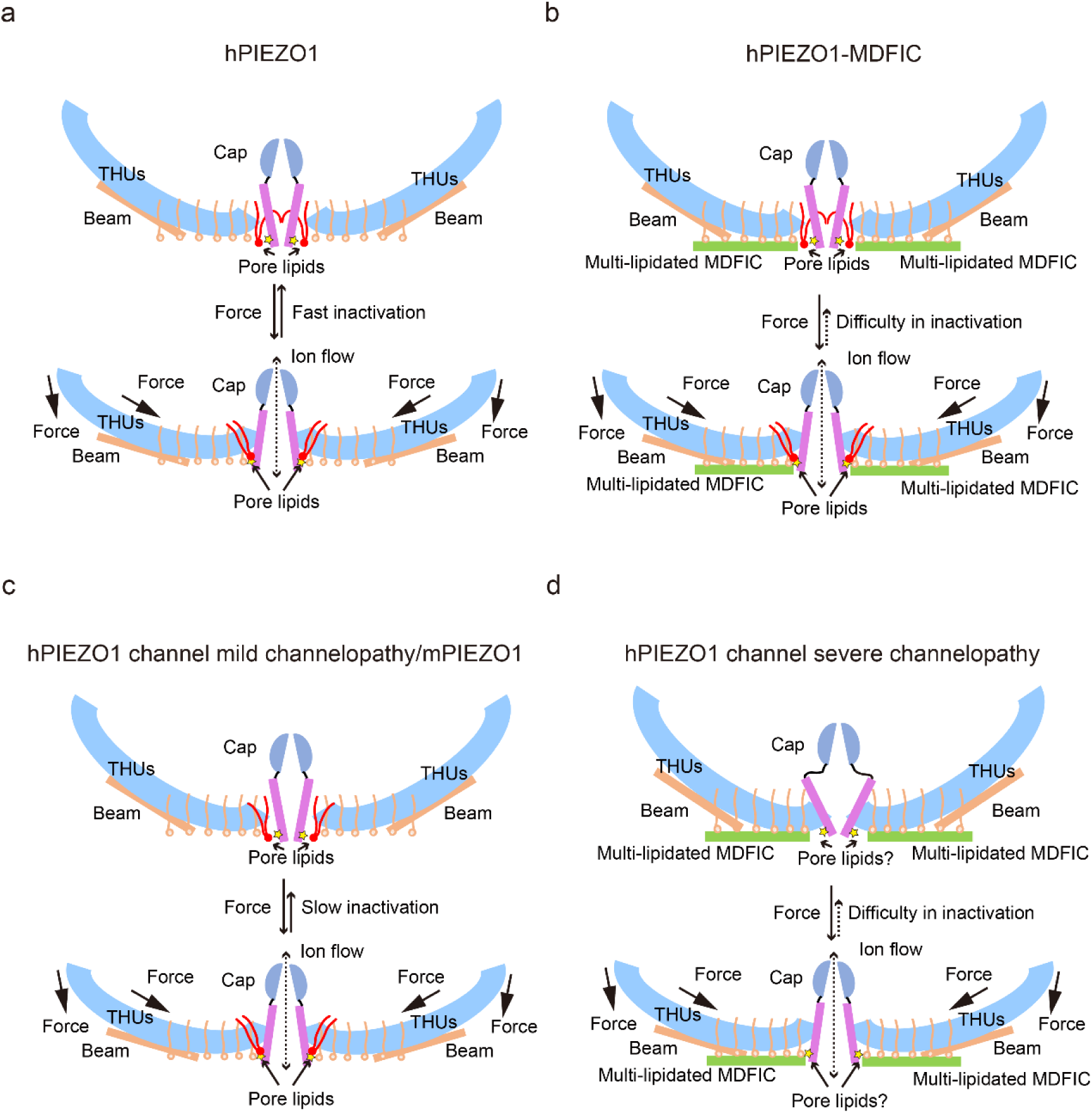
The proposed putative fast inactivation model of PIEZO1 channel. **a**, Cartoon representation of the fast inactivation model of the hPIEZO1 channel. Two of the three subunits are shown for clarity. The cap (purple semicircle), pore helices (pink rectangle), THUs (blue arc), beam (brown rectangle), and pore lipids (red) are present. The pore lipids interacting with R2456 (yellow star) seal the TM pore. The membrane force can reduce the membrane curvature and transduce the force to the pore region, removing the barrier created by the pore lipids, which might induce the channel pore to open. The pore lipids then rapidly return to the hydrophobic pore, exhibiting a fast-inactivation pattern. **b,** Cartoon diagram of the slow inactivation model of the hPIEZO1-MDFIC complex. Multi- lipidated MDFICs are shown as green rectangles. The fatty acid chains of the covalently linked MDFIC lipids can stabilize the pore lipids. Therefore, the hPIEZO1-MDFIC complex exhibits a very stable deep resting state. Once a higher threshold of mechanical force removes the pore lipids, it is also tricky for them to return to the hydrophobic pore region and reform the stable non- conducting pore module, thus exhibiting the prolonged slow inactivation phenotype. **c,** Cartoon diagram of the slow inactivation model of hPIEZO1 channel mild channelopathy or mPIEZO1. Mild channelopathy mutants or mPIEZO1 show the pore lipids retracted from the pore and thus exhibit a slow inactivation pattern. **d,** Cartoon representation of the slow inactivation model of the hPIEZO1 channel HX pathogenesis mutant R2456H with MDFIC auxiliary subunit. The R2456H channelopathy mutant probably disrupts the IH pore and the pore lipid interaction. Due to the stabilizing effect of the MDFIC auxiliary subunit, the pore has a twisted shape and the overall architecture of hPIEZO1-R2456H- MDFIC has a more curved and contracted blow-like shape.

The multi-lipidated MDFIC functions to stabilize the pore lipids that seal the hydrophobic pore further through interactions between hydrophobic acyl chain tails of the pore lipids and the lipids covalently linked to MDFIC (Fig. 7c and 7f). Once activated, the PIEZO-MDFIC complexes assume much slower inactivation (Fig. S2), consistent with the model that the multi-lipidated MDFIC makes the PIEZO challenging to open by mechanical force (Fig. 8b and 8d). Therefore, we deduce that the hPIEZO1-MDFIC/hPIEZO1-A1988V-MDFIC/hPIEZO1-E756del-MDFIC may reshape the pore module and fall into a deep resting state, in which it becomes rather tricky to remove the pore lipid from the hydrophobic pore by mild mechanical force. On the other hand, once the pore lipids are removed by a higher threshold of the mechanical force, it is equally more difficult for the pore lipids to return to the hydrophobic pore region and regain the stable pore module-multi-lipidated MDFIC- pore lipid complex, thus resulting in the very prolonged slow inactivation phenotype (Fig.8b and 8d).

## Discussion

Piezo ion channels are the critical force sensors^3^ that allow cells to sense their physical environment and regulate the cell fate. Intracellular signaling pathways have become the focus of research to understand the mechanism of mechanical force involved by piezo channels in cells^26^. MyoD was the first transcription factor identified to specify cell fate in a cell-autonomous fashion, ushering decades of investigation into cell fate control that led to the discovery of iPSC reprogramming^27^. Intriguingly, MyoD interacting proteins MDFIC and MDFI are auxiliary factors for the PIEZO mechanosensitive ion channels, suggesting that these complexes link the cell force sensing and cell fate control^19^. Indeed, mice with PIEZO deletions are lethal, supporting the idea that mechanosensing through these channels is critical in cell fate control during development. To understand the role of PIEZO or PIEZO-MDFIC/MDFI in human cell fate control, we solved the structure of hPIEZO1 in complex with and without MDFIC. MDFIC enables hPIEZO1 to respond to different forces by modifying the pore module through lipid interactions.

The fast and slow inactivation modes may allow cells to respond to external forces more accurately. Indeed, inactivation is widespread in different types of ion channels. Inactivation can be plastic, driven by intrinsic and extrinsic cues, and regulates many physiological processes. Mechanistically, inactivation follows different principles in different ion channels^28^. The inactivation rate of PIEZO channels is essential for the physiological functions of different cell types, including neuronal and non-neuronal cells. Moreover, different subtypes and species of PIEZO channels exhibit different inactivation rates. For example, hPIEZO1 and mPIEZO2 have a faster inactivation rate than mPIEZO1^3,9,18^. More importantly, abnormal inactivation of the PIEZO channel is one of the dominant outcomes of clinical PIEZO channelopathy^3,20^. The MDFIC inserts into the PIEZO pore module and significantly reshapes channel inactivation^19^, which may also link PIEZO channel inactivation to cell fate. The lack of knowledge regarding the faster inactivating hPIEZO1 has prevented the acquisition of structural information about the relationship of true fast inactivating wild-type hPIEZO1 between cell fate determination, as well as clinically significant hPIEZO1 GOF slow inactivating channelopathies. Our work has the following implications: First, we present the near-atomic cryo-EM structures of the fast inactivation hPIEZO1 and its slow inactivation channelopathy mutants, illuminating the fast inactivation mechanism of PIEZO channels involved by the pore lipids, which ingenious seals the hydrophobic pore.

Second, the overall structure of the curved hPIEZO1 shows a more flattened and extended state compared to the curved mPIEZO1, although there is no strong evidence yet for a link between curvature and inactivation. However, based on the mild GOF mutants hPIEZO1-A1988V and hPIEZO1- E756del, which are in the blade arm region, the correlation between curvature and inactivation is relatively evident, as the blade region indeed influences the channel inactivation. And all the force signal will also transduce to the pore region, removing the lipid barrier. Therefore, the pore lipid will likely play a key role in PIEZO gating.

Lastly, extrinsic cues, such as MDFIC, can modulate the inactivation, and the channelopathy mutants of PIEZO channels often cause a slower inactivation rate. The primary mechanism of PIEZO channel fast inactivation may provide clues against the clinical mechanopathologies. More importantly, these insights may inspire further investigation into mechanosensing channels as cell fate regulators in the future.

## Materials and Methods

### Constructs

A synthetic codon-optimized gene fragment encoding residues 1 to 2521 of the hPIEZO1 was cloned into a modified pEG-BacMam vector (Goehring et al., 2014) using EcoRI and XhoI restriction enzyme. The resulting protein has enhanced green fluorescent protein (EGFP) and a FLAG- modified antibody recognition sequence (DYKDDDDK) on the C terminus, separated by a PreScission protease (Ppase) cleavage site (LFQ/GP). Other mutations were built on its base by point mutations.

The cDNA for full-length human MDFIC residues 1–246) was fished from the human cDNA library and cloned into a similarly modified pEG-BacMam vector with no tag.

### Protein expression

Expi-HEK293F cells grown in HEK293 medium (Yocon) at 37°C with 6% CO2. *Spodoptera frugiperda* Sf9 cells were cultured in Sf-900 II SFM medium (Gibco) at 27°C. Cells were routinely tested for mycoplasma contamination and were negative.

hPIEZO1 and its mutations were expressed alone or co-expressed with the MDFIC subunit in expi- HEK293F cells using the BacMam technology. Bacmid carrying hPIEZO1 or MDFIC subunit was generated by transforming E.*coli* DH10Bac cells with the corresponding pEG-BacMam construct and Recombinant Bacmids were screened by blue-white spot validation. Baculoviruses were produced by transfecting Sf9 cells at a density of 1×10^6^ per ml with the bacmid using Cellfectin II (Invitrogen). To increase the viral titer, the recombinant virus has undergone two rounds of amplification to generate the P3 virus.

Expi-HEK293F cells in suspension were cultured to a density of 3×106 cells/ml and infected with the P3 virus. For expression of hPIEZO1 alone, cell culture was infected with 8% (v:v) of hPIEZO1 baculovirus. For co-expression of hPIEZO1 and MDFIC subunit, cell culture was infected with 4% (v:v) hPIEZO1 baculoviruses and 4% (v:v) MDFIC baculoviruses. After 12h, sodium butyrate was added to 10 mM final concentration and the temperature was decreased to 30°C. After 60 h of expression, cells were collected by centrifugation at 4000 r.p.m., 4°C for 10 min, resuspended in Tris-buffered saline (TBS) buffer containing 20 mM Tris pH 7.4, 150 mM NaCl.

### Protein purification

The cell pellet was homogenized in a TBS buffer supplemented with 2 mM phenylmethylsulfonyl fluoride (PMSF) by ultrasonication. Then, large organelles and insoluble matter were pelleted by centrifugation at 8000 g for 10 min. The supernatant was centrifuged at 36,000 r.p.m for 30 min in a Ti45 rotor (Beckman). The membrane pellet was mechanically homogenized and solubilized in extraction buffer containing TBS, 1% (w/v) lauryl maltose neopentyl glycol (LMNG) and 0.1% (w/v) cholesteryl hemisuccinate (CHS) for an hour with stirring. Insoluble materials were removed by centrifugation at 36,000 r.p.m. for 1 h in a Ti45 rotor (Beckman). The supernatant was loaded onto anti-FLAG G1 affinity resin (GenScript) by gravity flow. The resin was further washed with 10 column volumes of wash buffer containing TBS and 0.02% (w/v) LMNG, and protein was eluted with an elution buffer containing TBS, 0.02% (w/v) LMNG, and 230 μg/ml FLAG peptide. The C-terminal EGFP tag of eluted protein was removed by Ppase cleavage at 4 °C overnight. The protein was further concentrated by a 100-kDa cutoff concentrator (Millipore) and loaded onto a Superose 6 increase 10/300 column (GE Healthcare) running in TBS with 0.01% (w/v) digitonin. Peak fractions were combined for cryo-EM sample preparation.

### Cryo-EM sample preparation

For cryo-EM sample preparation, 3 μl aliquots of the protein sample were loaded onto glow-discharged (20 s, 15 mA; Pelco easiGlow, Ted Pella) Au grids (Quantifoil, Au R1.2/1.3, 300 mesh). The grids were blotted for 6 s with 3 forces after waiting for 20 s and immersed in liquid ethane using Vitrobot (Mark IV, Thermo Fisher Scientific) in 100% humidity and 8°C.

### Data collection

Cryo-EM data were collected at a nominal magnification of 215K (resulting in a calibrated pixel size of 0.57 Å) on a Titan Krios (Thermo Fisher Scientific) operating at 300 kV equipped with a K3 or Falcon4i Summit detector and GIF Quantum energy filter (slit width 20-eV) in super-resolution mode. Movie stacks were automatically acquired using EPU software. The defocus range was set from −0.9 to −1.3 μm. Each movie stack, consisting of 32 frames, was exposed for 2.72 seconds with a total dose of ∼40 e^−^/Å^2^.

### Image processing and model building

Data processing was carried out with cryoSPARC suite^29^. Patch CTF estimation was carried out after alignment and summary of all 32 frames in each stack using the patch motion correction. Initial particles were picked from a few micrographs using blob picker in cryoSPARC and 2D averages were generated. Final particle picking was done by template picker using templates from those 2D results. After three rounds of 2D classification, ab-initio reconstruction, non-uniform refinement and local refinement for reconstructing the density map. All maps were low-pass filtered to the map-model FSC value. The reported resolutions were based on the FSC=0.143 criterion. An initial model was generated by mPIEZO1 (7WLT). Then, we manually completed and refined the model using Coot. Subsequently, the models were refined against the corresponding maps by PHENIX. We used PyMol and UCSF Chimera^30^ for structural analysis and graphics generation.

### Electrophysiological recording

PIEZO1-KO-HEK293T cells were a gift obtained from Bailong Xiao Lab and cultured on coverslips placed in a 12-well plate containing DMEM medium (Gibco) supplemented with 10% fetal bovine serum (FBS). The cells in each well were transiently transfected with 1 μg hPIEZO1 or mutant hPIEZO1 plasmids fused with GFP or co-transfected with MDFIC plasmid (weigh ratio=1:3) using polyethyleneimine (PEI) according to the manufacturer’s instructions. After 12–20 h, the coverslips were transferred to a recording chamber containing the external solution (10 mM HEPES-Na pH 7.4, 150 mM NaCl, 5 mM glucose, 2 mM MgCl2, and 1 mM CaCl2). Borosilicate micropipettes (OD 1.5 mm, ID 0.86 mm, Sutter) were pulled and fire-polished to 2–5 MΩ resistance. For whole-cell recordings, the pipette solution was 10 mM HEPES-Na pH 7.4, 150 mM CsCl, and 5 mM EGTA. The bath solution was 10 mM HEPES-Na pH 7.4, 150 mM NaCl, 5 mM glucose, 2 mM MgCl2, and 1 mM CaCl2.

Recordings were obtained at room temperature (∼25 °C) using an Axopatch 200B amplifier, a Digidata 1550 digitizer, and pCLAMP 10.7 software (Molecular Devices). The patches were held at −80 mV and the recordings were low-pass filtered at 1 kHz and sampled at 20 kHz. Mechanical poking was delivered to the cell being patched under whole-cell configuration at an angle of 80° using a fire-polished glass pipette (the tip diameter 3-4 mm). Downward movement of the probe toward the cell was driven by a Clampex-controlled piezo-electric crystal micro-stage (Physik Instrument; E625 LVPZT Controller/Amplifier). The probe had a velocity of 1 μm/ms during the downward/upward motion and the stimulus was maintained for 1 s. A series of mechanical steps in 1 mm increments was applied every 1.4 s.

### Reporting summary

Further information on research design is available in the Nature Portfolio Reporting Summary linked to this article.

### Data availability

Cryo-EM maps and related structure coordinates of hPIEZO1, hPIEZO1-MDFIC, hPIEZO1- A1988V, hPIEZO1-A1988V-MDFIC, hPIEZO1-E756del-MDFIC and hPIEZO1-R2456H-MDFIC have been deposited in the EMDB and PDB under accession codes EMD-39205 (PDB 8YEZ), EMD-60479 (PDB 8ZU3), EMD-60481 (PDB 8ZU8), EMD-39219 (PDB 8YFC), EMD-39222 (PDB 8YFF), and EMD-39223 (PDB 8YFG), respectively. Source data are provided in this paper.

## Acknowledgments

We want to thank the Cryo-EM Facility and High-Performance Computing (HPC) Center of Westlake University for providing cryo-EM and computation support. This work was supported by National Natural Science Foundation of China (92068201) and Key R&D Program of Zhejiang (2024SSYS0031). We also would like to thank all the Cell fate control lab members for their support.

## Author contributions

M.Z. conceived the project and designed the experiments. Y.S. and X.G. prepared the constructs and purified the proteins. Y.S., M.Z. and Y.L. prepared the cryo-EM sample and collected cryo-EM data. M.C. performed the electrophysiological study. M.Z. and Y.S. performed image processing, built the model, analyzed data, and wrote the manuscript draft. D.P. and M.Z. supervised the project. All authors contributed to the manuscript preparation.

---

**Figure. S1.**
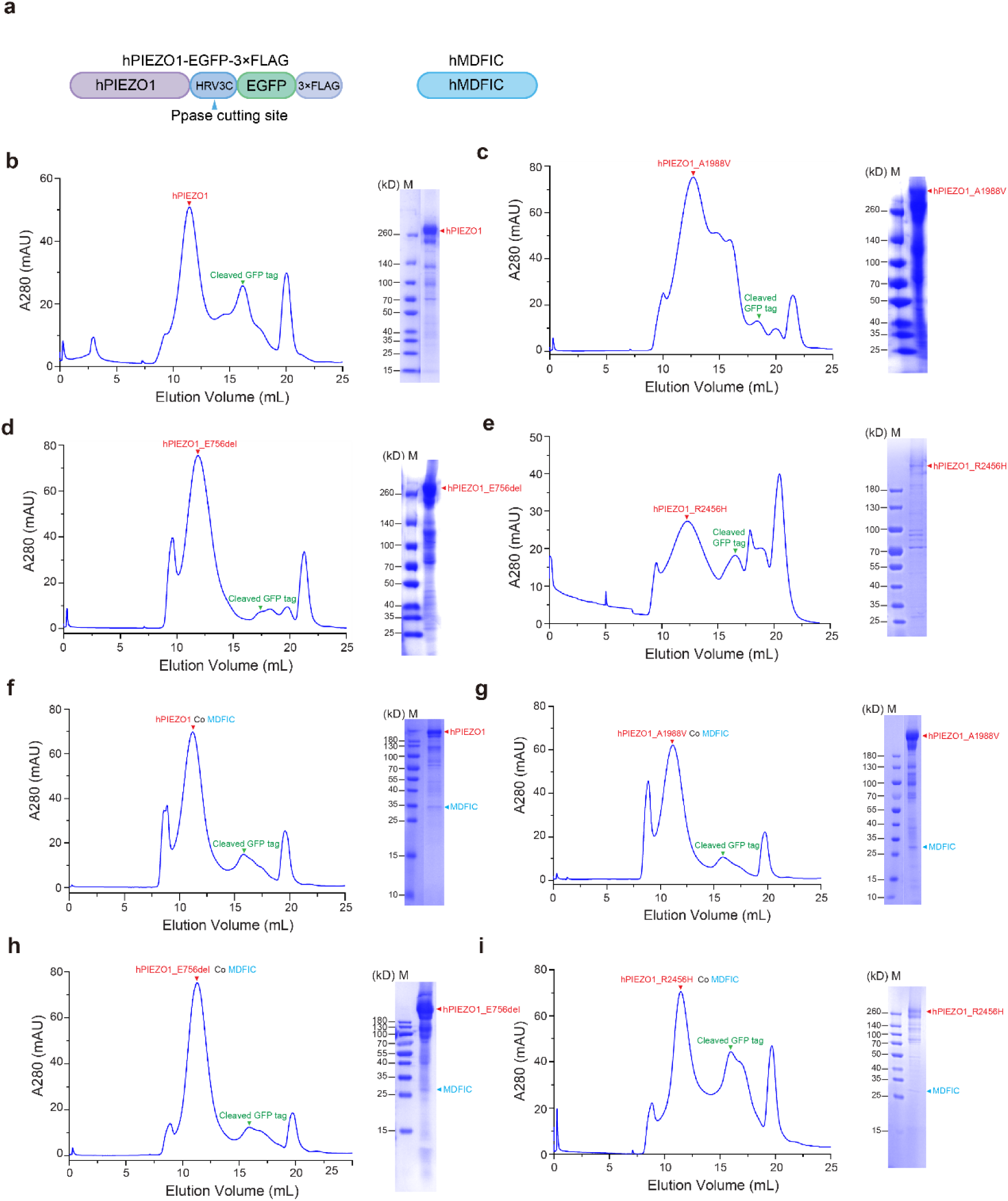
Purification procedures of hPIEZO1, the channelopathy mutants with or without MDFIC proteins. **a**, A schematic diagram of the hPIEZO1-GFP-3×FLAG constructs for protein purifications and electrophysiological studies. **b-e**, Representative size-exclusion chromatography (SEC) trances and Coomassie-staining SDS- PAGE analysis of purified WT hPIEZO1 (b), hPIEZO1-R2456H (c), hPIEZO1-A1988V (d) and hPIEZO1-E756del (e) protein samples. Red arrows denote the peak position and bands of hPIEZO1 and its mutant. Green arrows denote the peak position of the cleaved GFP tag. **f-g**, Representative size-exclusion chromatography (SEC) trances and Coomassie-staining SDS- PAGE analysis of purified WT hPIEZO1 co hMDFIC (b), hPIEZO1-R2456H co hMDFIC (c), hPIEZO1-A1988V co hMDFIC (d) and hPIEZO1-E756del co hMDFIC (e) protein samples. The peak positions and bands of hPIEZO1 and its mutant are denoted by red arrows. Green arrows indicate the peak position of the cleaved GFP tag. Blue arrows represent the bands of hMDFIC.

**Figure. S2.**
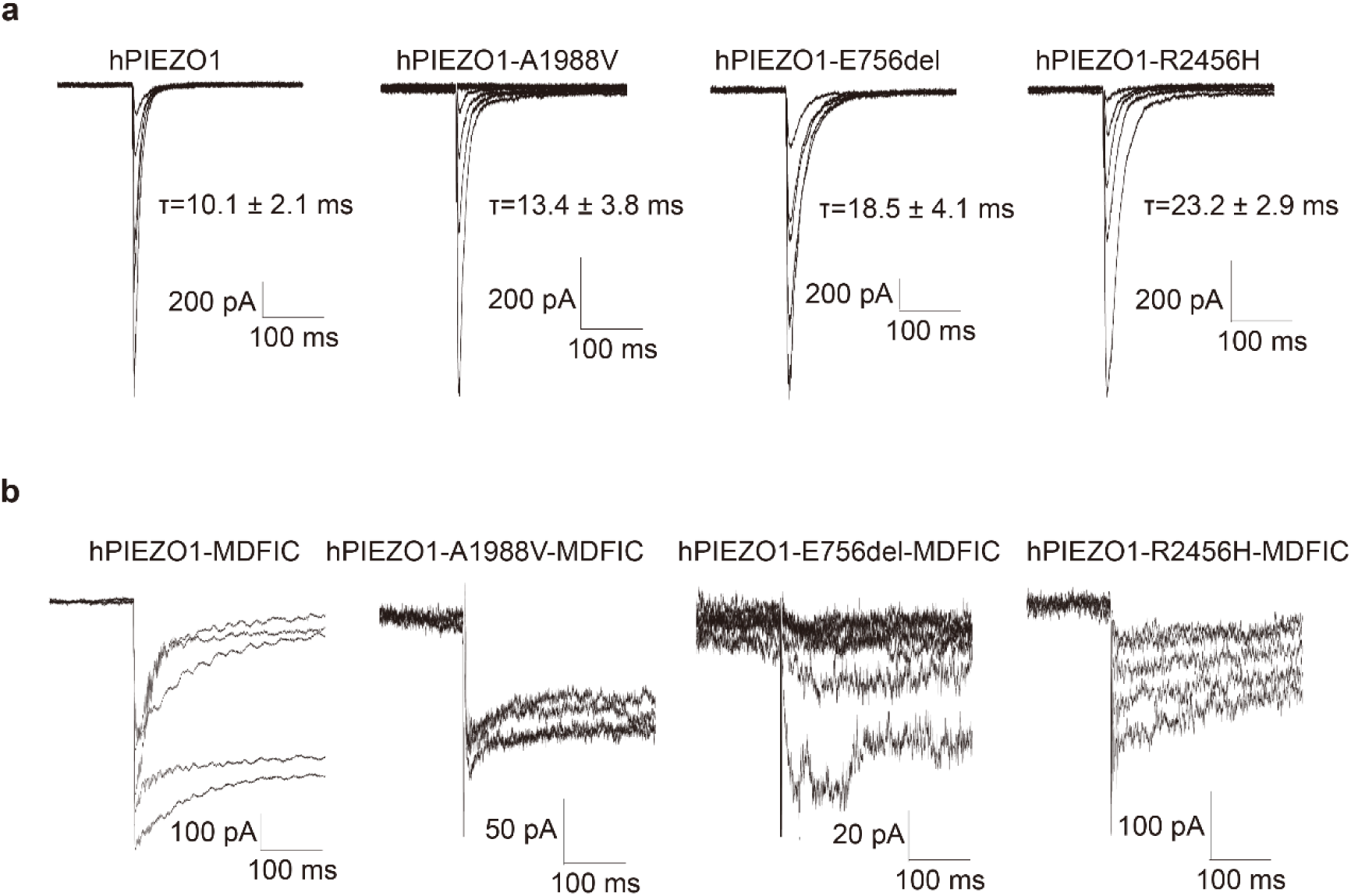
The whole-cell poking evoked currents of wild-type hPIEZO1, the GOF channelopathy mutants with or without MDFIC. **a**, The representative traces of whole-cell poking evoked currents of wild-type hPIEZO1, hPIEZO1- A1988V, hPIEZO1-E756del and hPIEZO1-R2456H subjected to a series of mechanical steps in 1 µm increments from 5 μm at a holding potential of −60 mV in HEK293T-PIEZO1-KO cells. The hPIEZO1 shows a very fast inactivation behavior with a half inactivation time of 10.1 ± 2.1 ms. At the same time, the channelopathy mutants A1988V, E756del and R2456H present a slow inactivation rate with half inactivation time of 13.4 ± 3.8 ms, 18.5 ± 4.1 ms and 23.2 ± 2.9 ms, respectively. **b,** The representative traces of whole-cell poking evoked currents of wild-type hPIEZO1, hPIEZO1- A1988V, hPIEZO1-E756del and hPIEZO1-R2456H co-expressed with the auxiliary subunit MDFIC subjected to a series of mechanical steps in 1 µm increments from 8 μm at a holding potential of −60 mV in HEK293T-PIEZO1-KO cells. The evoking MS currents were negligible even with large increments, presenting the significant slow inactivation behavior. Therefore, the MDFIC may attenuate the mechanosensitivity of wild-type hPIEZO1 and GOF channelopathy mutants. All data are replicated at least three times and the statistic is based on the mean ± SEM.

**Figure. S3.**
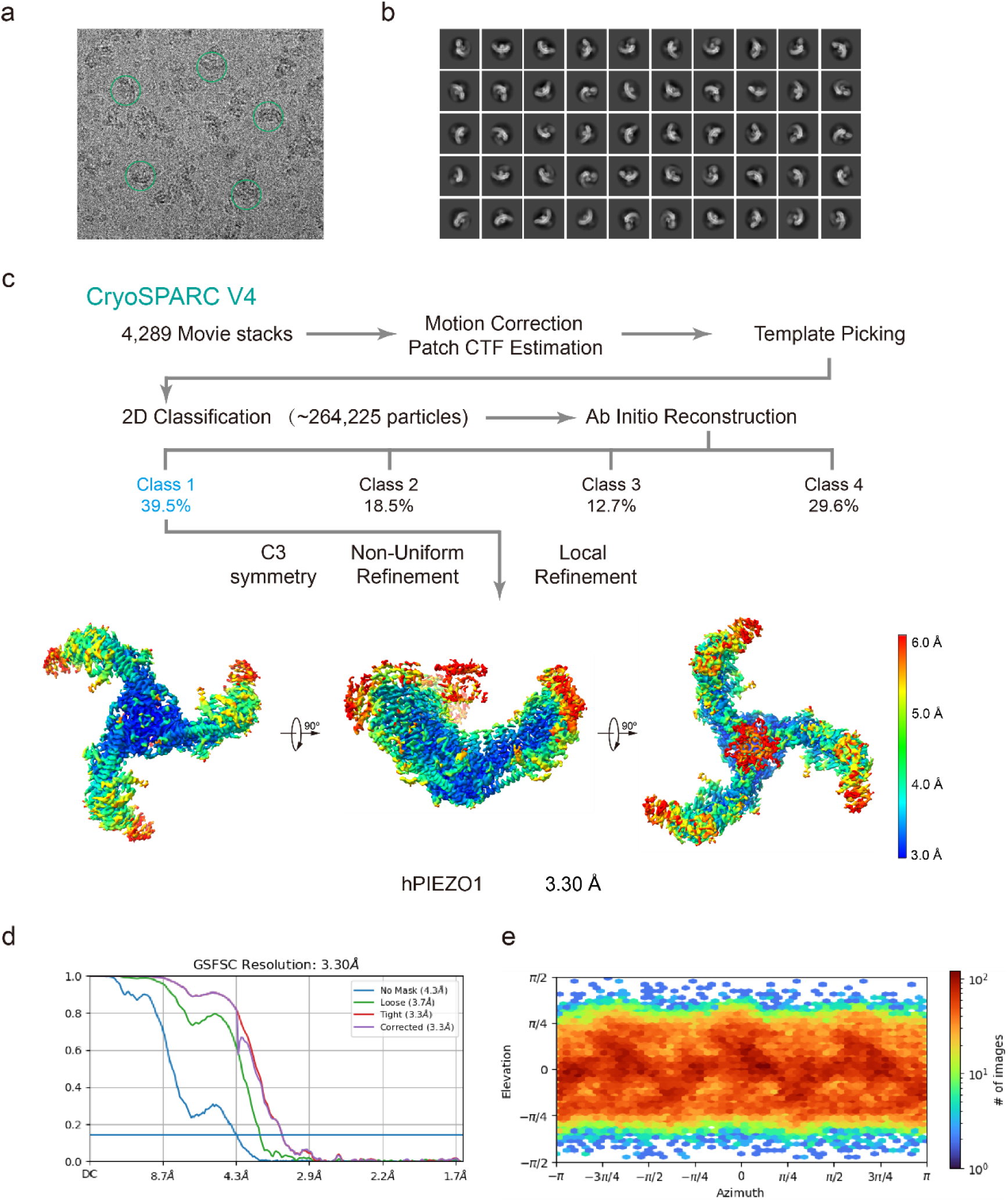
Cryo-EM data processing procedure of WT hPIEZO1. **a**, Representative raw micrograph of hPIEZO1. **b**, Representative 2D class averages of the cryo-EM particles of hPIEZO1. **c**, Flowchart of the image processing procedure for hPIEZO1. Resolution estimation for the tight masked map is based on the criterion of an FSC cutoff of 0.143. **d-e**, Gold-standard Fourier shell correlation (FSC) curves of the final refined maps (d) and angular distribution histogram (e) of the final hPIEZO1 reconstruction. This is a standard output from cryoSPARC v4.

**Figure. S4.**
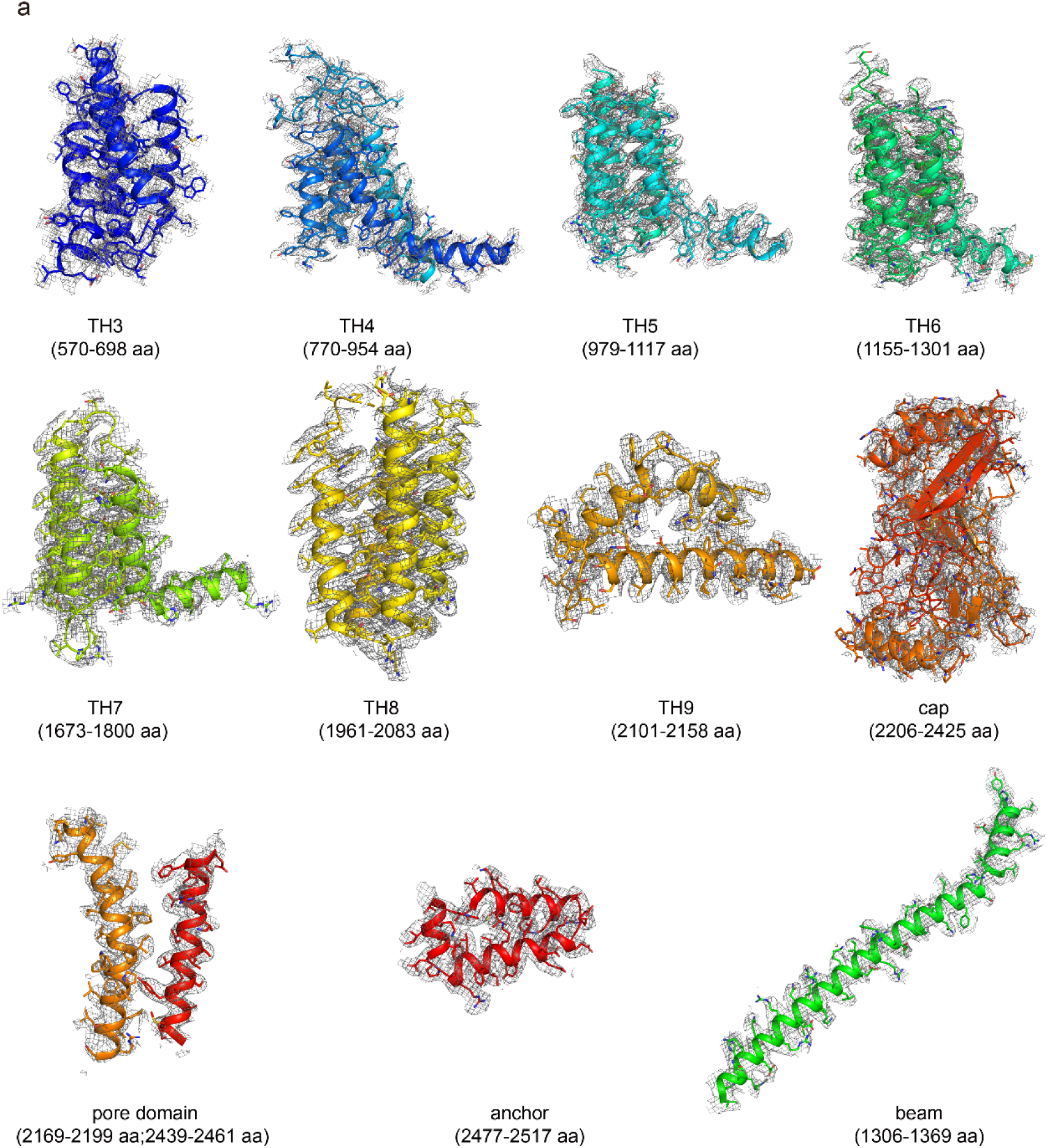
The EM density of WT hPIEZO1. The EM density segments of the transmembrane helices domain (TH3-TH9), cap domain, pore domain, anchor and beam of WT hPIEZO1.

**Figure. S5.**
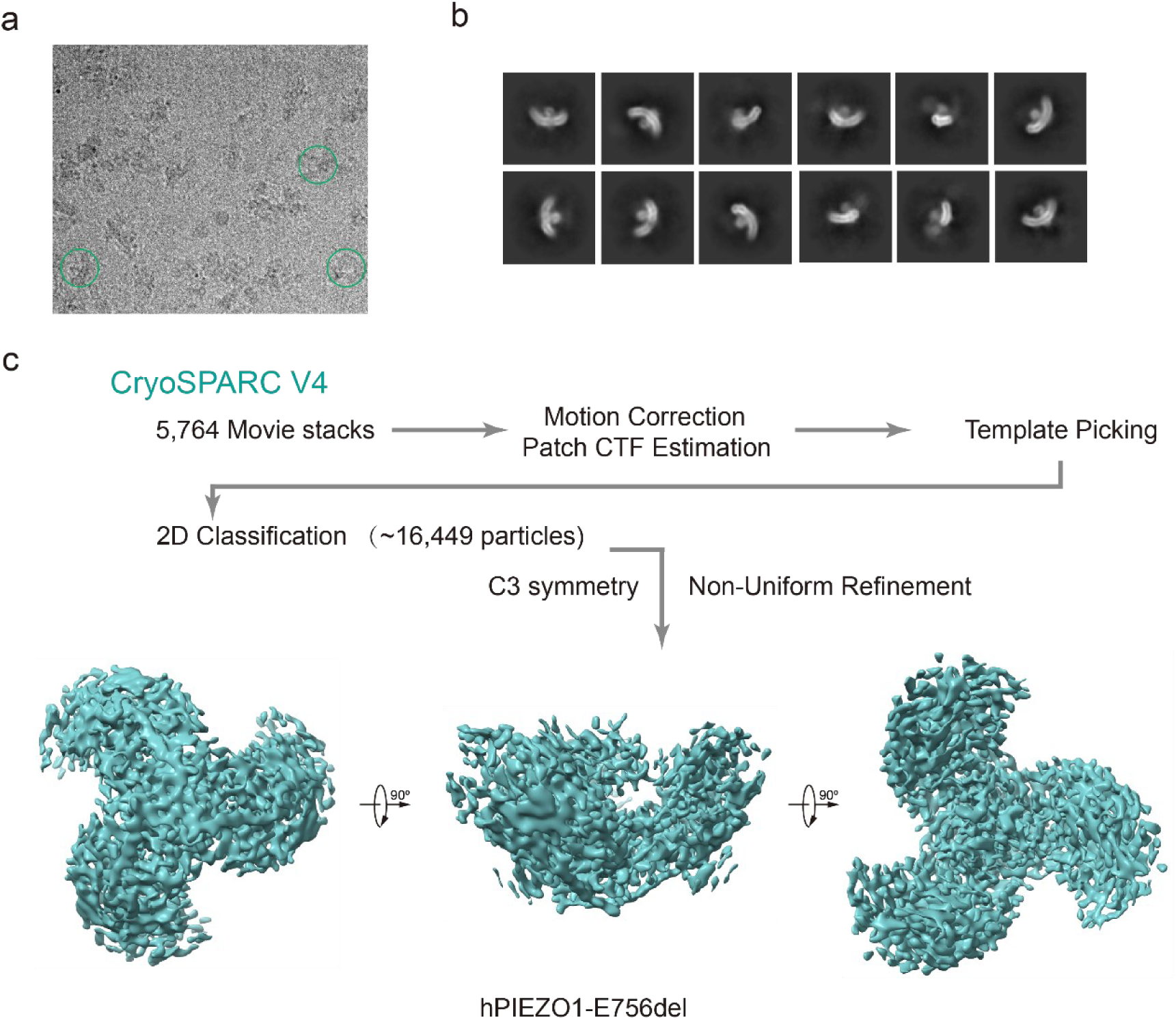
Cryo-EM data processing procedure of hPIEZO1-E756del mutant. **a**, Representative raw micrograph of hPIEZO1-E756del. **b**, Representative 2D class averages of the cryo-EM particles of hPIEZO1- E756del. **c**, Flowchart of the image processing procedure for hPIEZO1- E756del. This is a standard output from cryoSPARC v4.

**Figure. S6.**
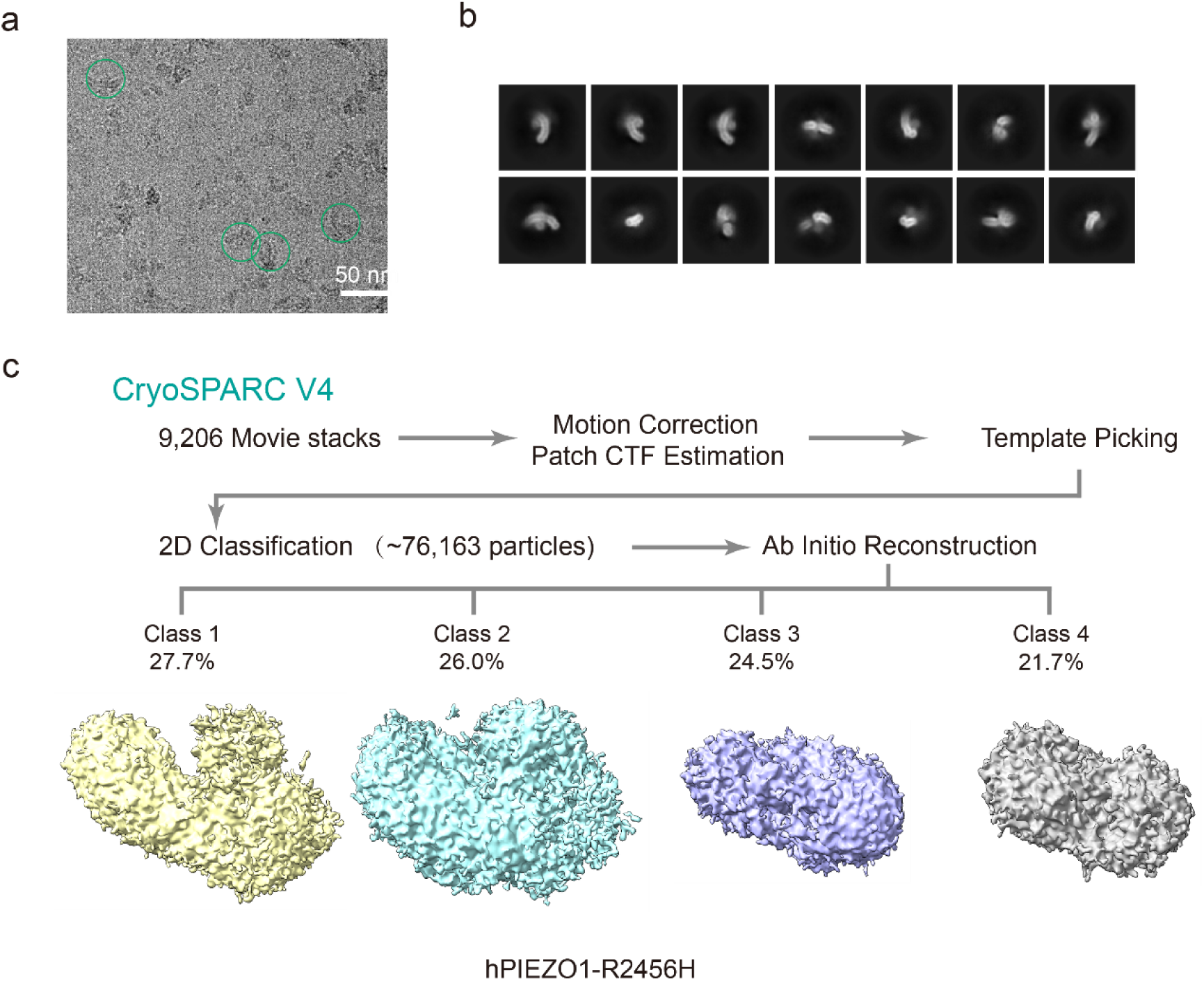
Cryo-EM data processing procedure of hPIEZO1-R2456H mutant. **a**, Representative raw micrograph of hPIEZO1-R2456H. **b**, Representative 2D class averages of the cryo-EM particles of hPIEZO1-R2456H. **c**, Flowchart of the image processing procedure for hPIEZO1-R2456H. This is a standard output from cryoSPARC v4.

**Extended Data Fig. 7.**
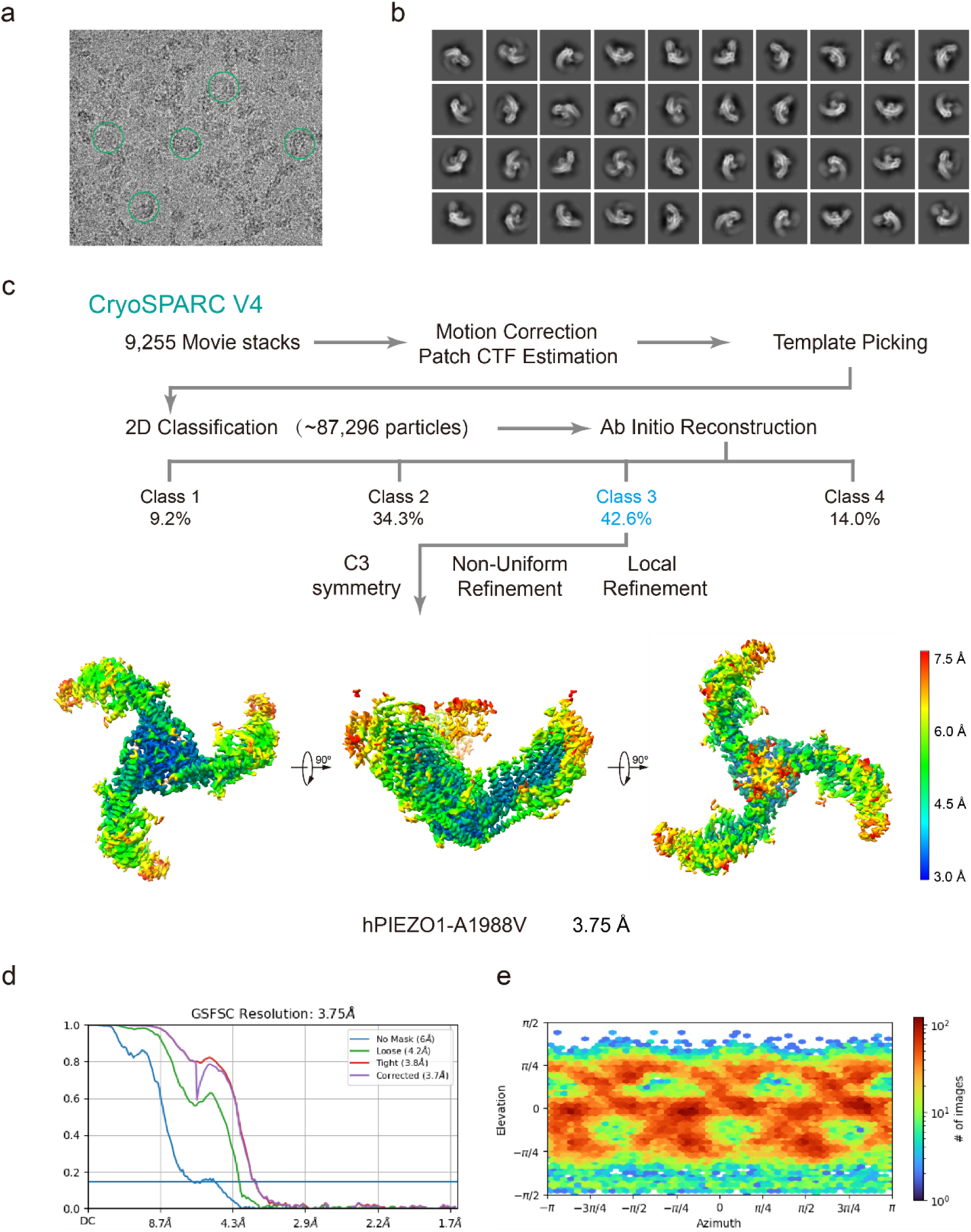
Cryo-EM data processing procedure of hPIEZO1-A1988V mutant. a, Representative raw micrograph of hPIEZO1-A1988V. **b**, Representative 2D class averages of the cryo-EM particles of hPIEZO1- A1988V. **c**, Flowchart of the image processing procedure for hPIEZO1- A1988V. Resolution estimation for the tight masked map is based on the criterion of an FSC cutoff of 0.143. **d-e**, Gold-standard Fourier shell correlation (FSC) curves of the final refined maps (d) and angular distribution histogram (e) of the final hPIEZO1- A1988V reconstruction. This is a standard output from cryoSPARC v4.

**Extended Data Fig. 8.**
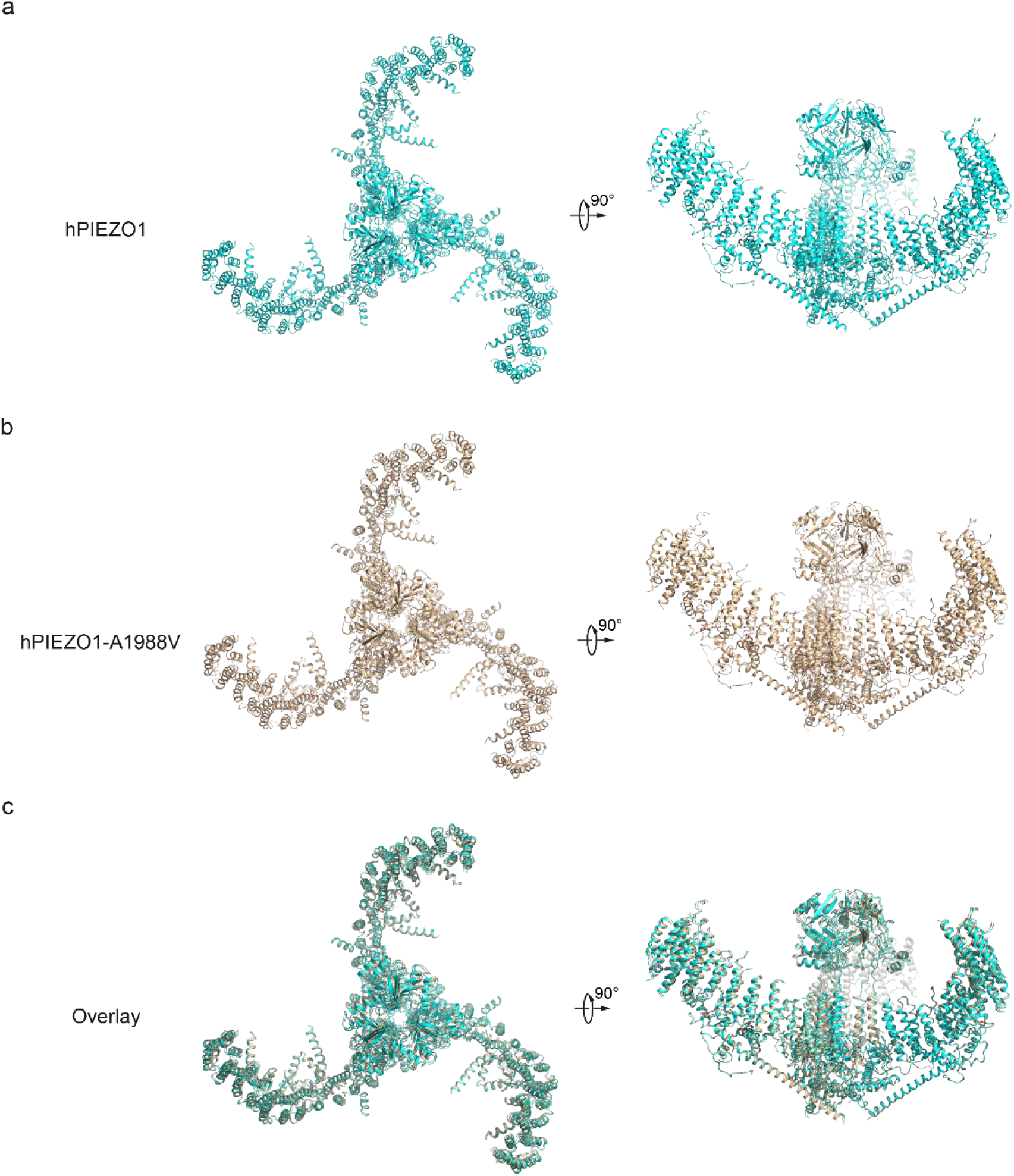
Structural comparison of hPIEZO1 and hPIEZO1-A1988V a, Overall structure of hPIEZO1 viewed from the top (left) and side (right). **b**, Overall structure of hPIEZO1-A1998V viewed from the top (left) and side (right). **c**, Overall structure comparison of hPIEZO1 (cyan) and hPIEZO1-A1998V (brown) viewed from the top (left) and side (right).

**Extended Data Fig. 9.**
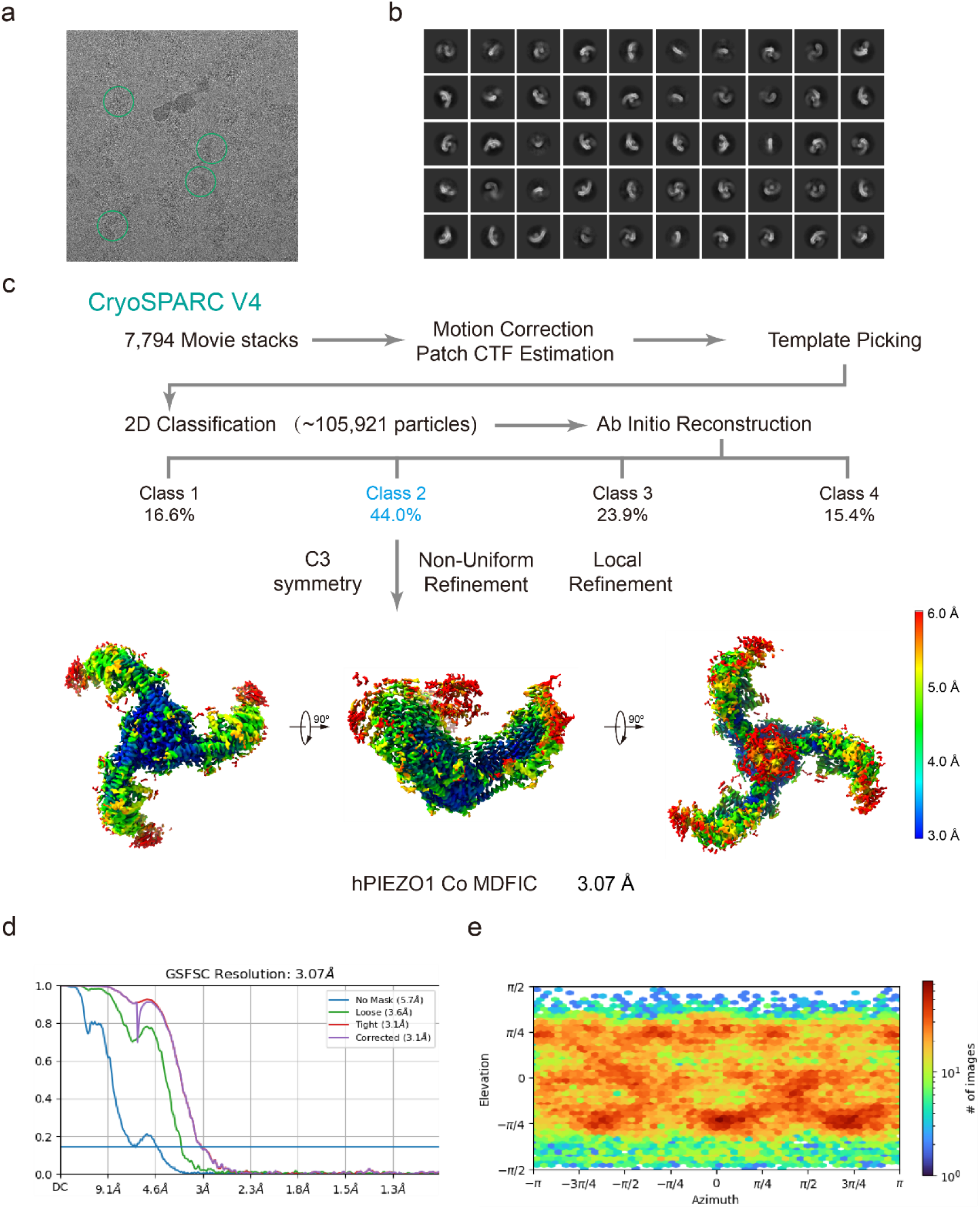
Cryo-EM data processing procedure of hPIEZO1 Co MDFIC. **a**, Representative raw micrograph of hPIEZO1 co MDFIC. **b**, Representative 2D class averages of the cryo-EM particles of hPIEZO1 co MDFIC. **c**, Flowchart of the image processing procedure for hPIEZO1 co MDFIC. Resolution estimation for the tight masked map is based on the criterion of an FSC cutoff of 0.143. **d-e**, Gold-standard Fourier shell correlation (FSC) curves of the final refined maps (d) and angular distribution histogram (e) of the final hPIEZO1 co MDFIC reconstruction. This is a standard output from cryoSPARC v4.

**Extended Data Fig. 10.**
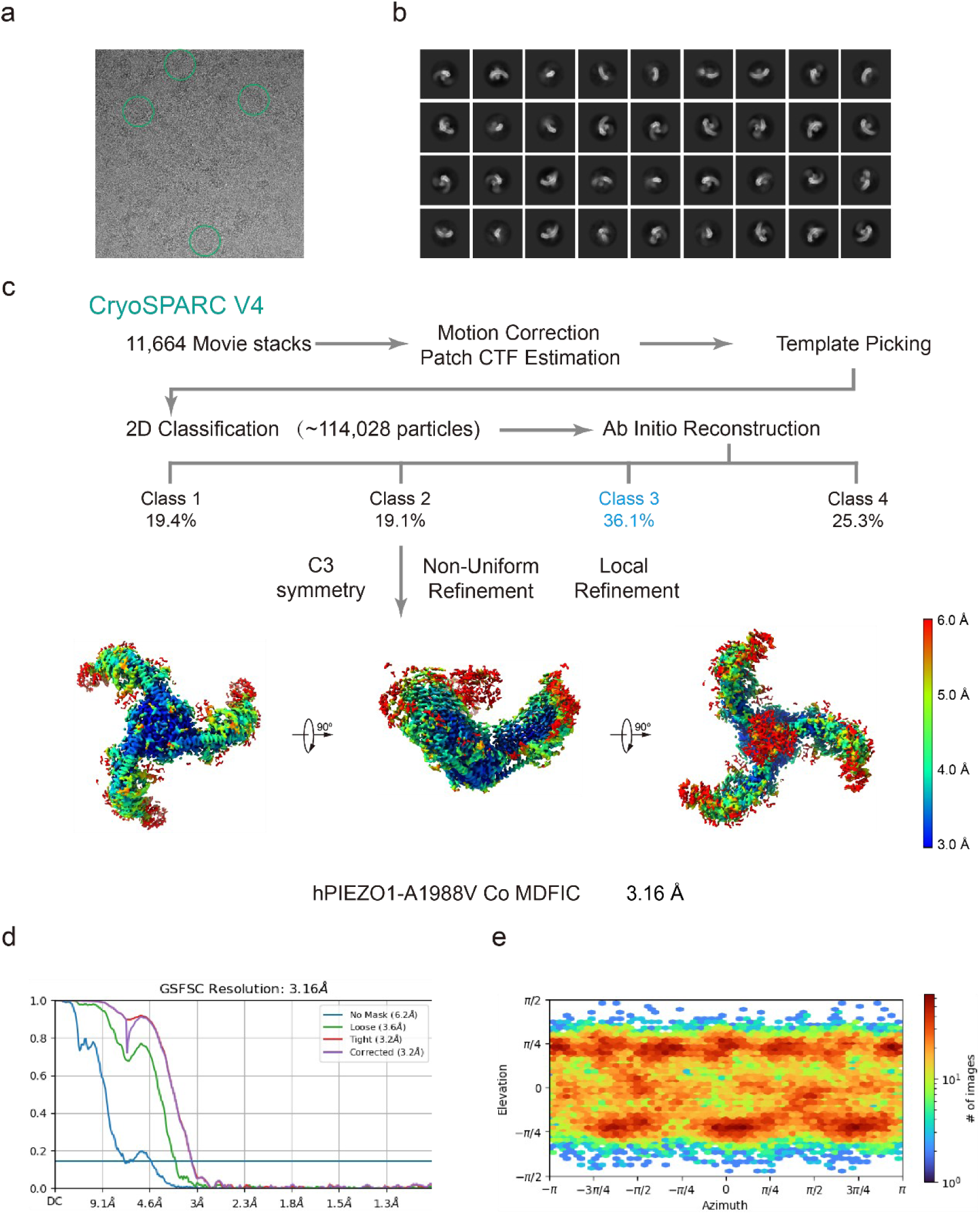
Cryo-EM data processing procedure of hPIEZO1-A1988V Co MDFIC. a, Representative raw micrograph of hPIEZO1-A1988V co MDFIC. **b**, Representative 2D class averages of the cryo-EM particles of hPIEZO1-A1988V co MDFIC. **c**, Flowchart of the image processing procedure for hPIEZO1-A1988V co MDFIC. Resolution estimation for the tight masked map is based on the criterion of an FSC cutoff of 0.143. **d-e**, Gold-standard Fourier shell correlation (FSC) curves of the final refined maps (d) and angular distribution histogram (e) of the final hPIEZO1-A1988V co MDFIC reconstruction. This is a standard output from cryoSPARC v4.

**Extended Data Fig. 11.**
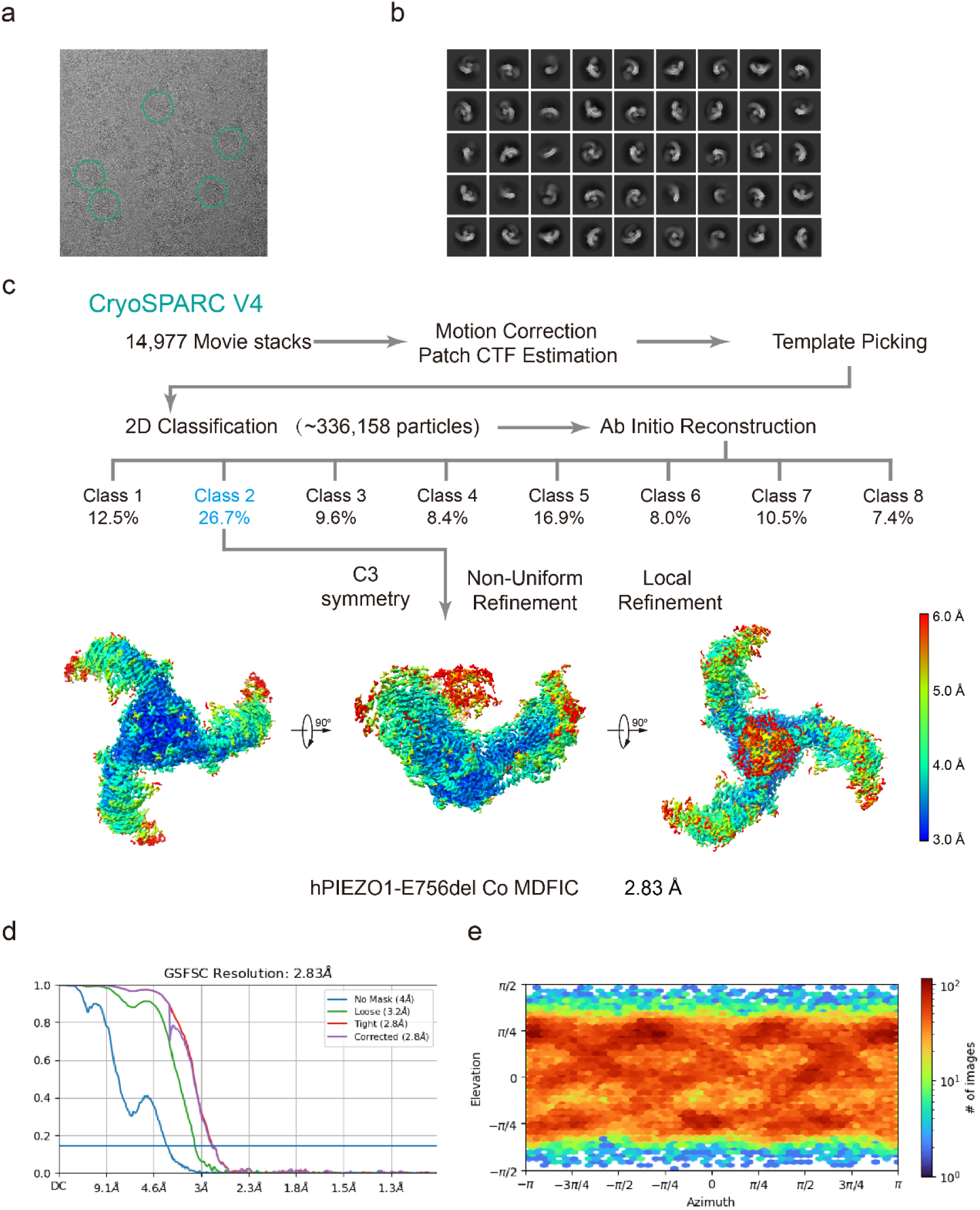
Cryo-EM data processing procedure of hPIEZO1-E756del Co MDFIC. **a**, Representative raw micrograph of hPIEZO1-E756del co MDFIC. **b**, Representative 2D class averages of the cryo-EM particles of hPIEZO1-E756del co MDFIC. **c**, Flowchart of the image processing procedure for hPIEZO1-E756del co MDFIC. Resolution estimation for the tight masked map is based on the criterion of an FSC cutoff of 0.143. **d-e**, Gold-standard Fourier shell correlation (FSC) curves of the final refined maps (d) and angular distribution histogram (e) of the final hPIEZO1-E756del co MDFIC reconstruction. This is a standard output from cryoSPARC v4.

**Extended Data Fig. 12.**
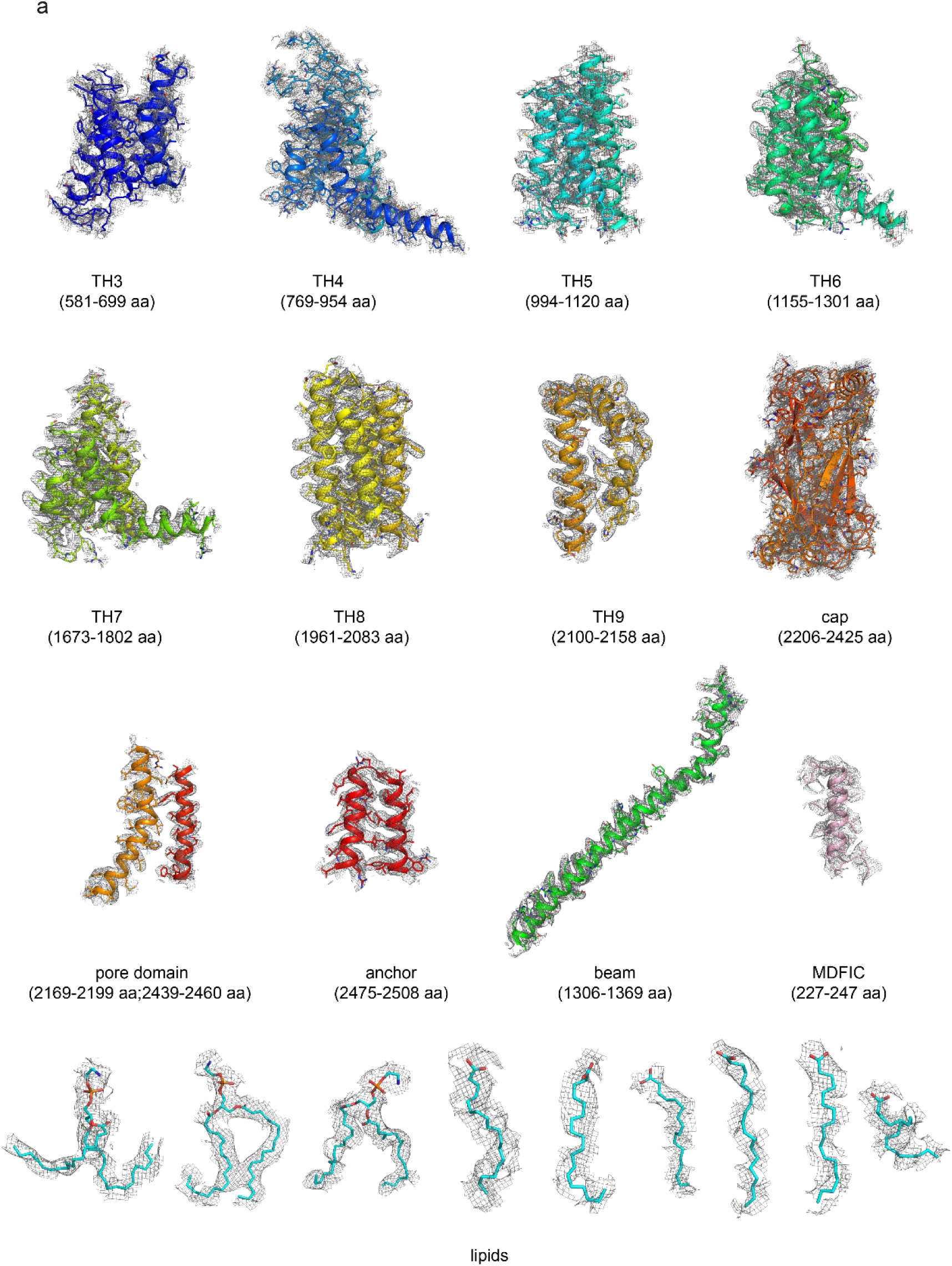
The EM density of hPIEZO1-E756del-MDFIC. The EM density segments of the transmembrane helices domain (TH3-TH9), cap domain, pore domain, anchor and beam of hPIEZO1-E756del. The EM density of the auxiliary subunit MDFIC and the bound lipids is also shown.

**Extended Data Fig. 13.**
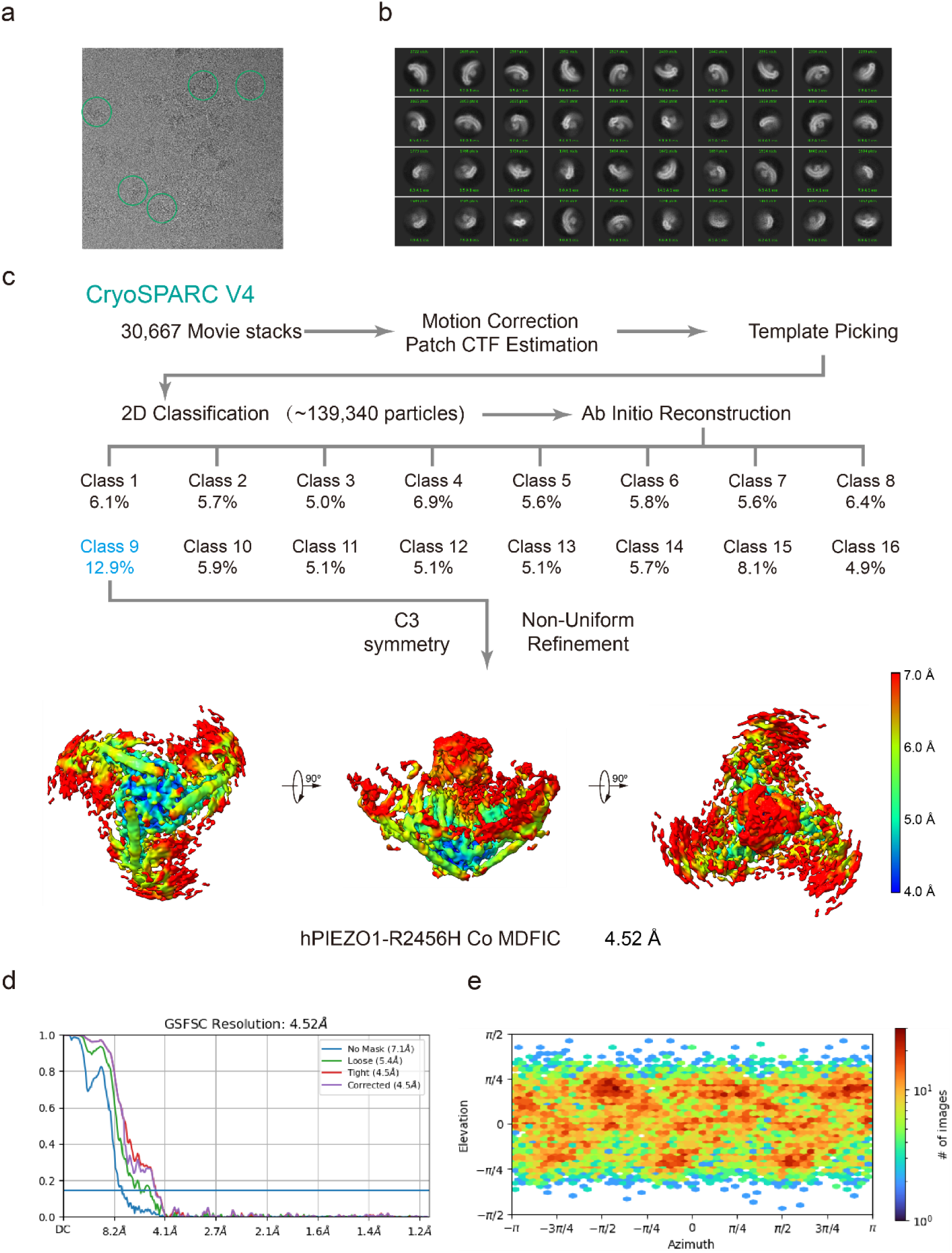
Cryo-EM data processing procedure of hPIEZO1-R2456H Co MDFIC. a, Representative raw micrograph of hPIEZO1-R2456H co MDFIC. **b**, Representative 2D class averages of the cryo-EM particles of hPIEZO1-R2456H co MDFIC. **c**, Flowchart of the image processing procedure for hPIEZO1-R2456H co MDFIC. Resolution estimation for the tight masked map is based on the criterion of an FSC cutoff of 0.143. **d-e**, Gold-standard Fourier shell correlation (FSC) curves of the final refined maps (d) and angular distribution histogram (e) of the final hPIEZO1-R2456H co MDFIC reconstruction. This is a standard output from cryoSPARC v4.

**Extended Data Fig. 14.**
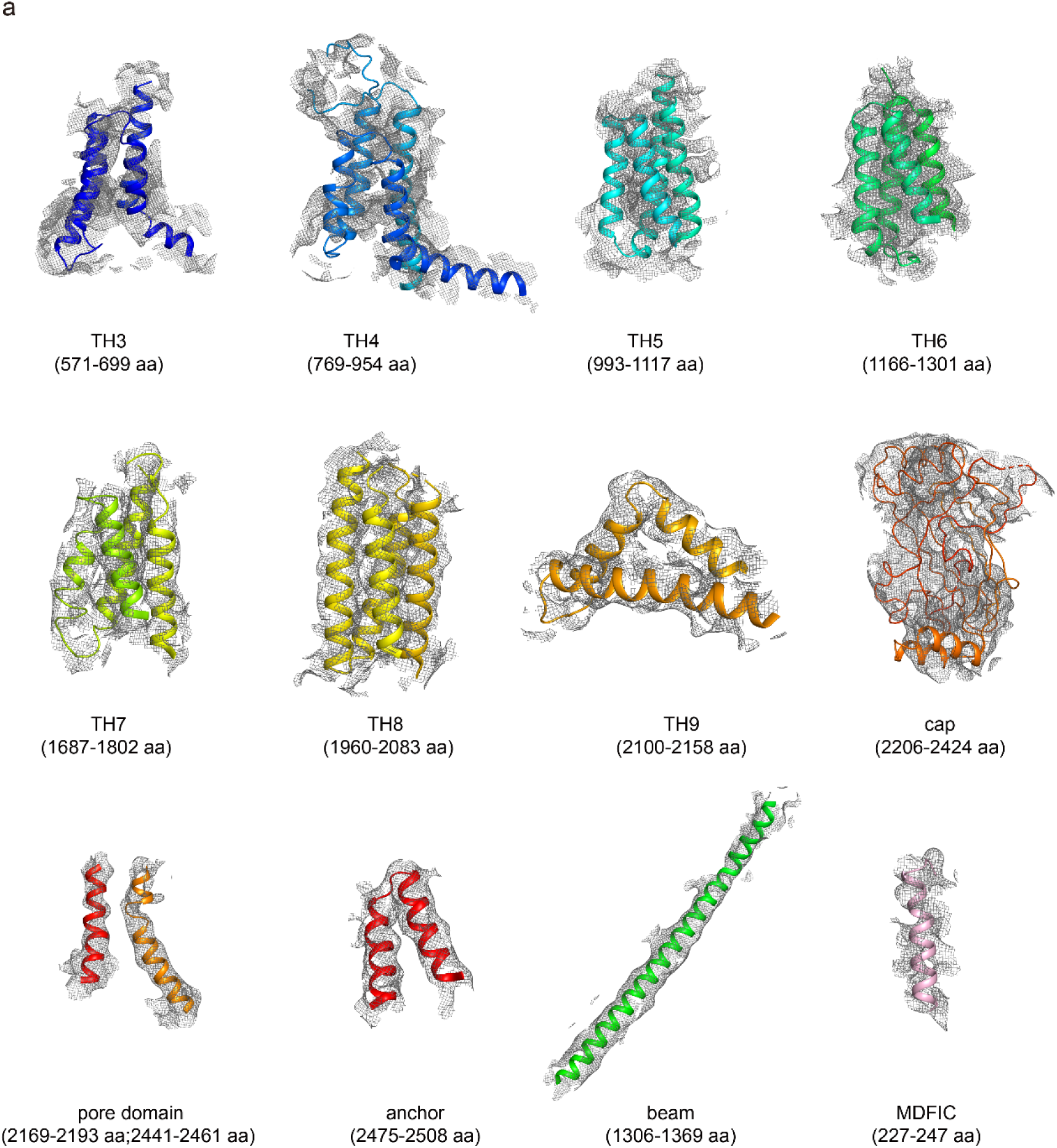
The EM density of hPIEZO1-R2456H Co MDFIC. The EM density segments of the transmembrane helices domain (TH3-TH9), cap domain, pore domain, anchor and beam of hPIEZO1-R2456H Co MDFIC. The EM density of the auxiliary subunit MDFIC and the bound lipids is also shown.

**Extended Data Table 1.** Cryo-EM data collection, refinement, and validation statistics.

|  | hPIEZO1 | hPIEZO1-A1988V | hPIEZO1 Co<br>MDFIC | hPIEZO1-A1988V Co<br>MDFIC | hPEIZO1-E756del Co<br>MDFIC | hPEIZO1-R2456H Co<br>MDFIC |
| --- | --- | --- | --- | --- | --- | --- |
| **Data collection and processing** |  |  |  |  |  |  |
| Microscope | FEI Titan | FEI Titan | FEI Titan | FEI Titan | FEI Titan | FEI Titan |
|  | Krios | Krios | Krios | Krios | Krios | Krios |
| Magnification | 105,000 | 215,000 | 215,000 | 215,000 | 215,000 | 215,000 |
| Voltage (KV) | 300 | 300 | 300 | 300 | 300 | 300 |
| Detector | Gatan K3 | Falcon 4i | Falcon 4i | Falcon 4i | Falcon 4i | Falcon 4i |
| Electron exposure (e<sup>-</sup>/ Å<sup>2</sup>) | 60 | 60 | 40 | 40 | 40 | 40 |
| Defocus range (μm) | -0.9 to -1.3 | -0.9 to -1.3 | -0.9 to -1.3 | -0.9 to -1.3 | -0.9 to -1.3 | -0.9 to -1.3 |
| Pixel size (Å) | 0.849 | 0.849 | 0.57 | 0.57 | 0.57 | 0.57 |
| Symmetry imposed | C3 | C3 | C3 | C3 | C3 | C3 |
| Initial particle images (no.) | 264,225 | 87,296 | 105,921 | 114,028 | 336,158 | 139,340 |
| Final particle images (no.) | 104,369 | 37,188 | 46,605 | 41,164 | 89,754 | 17,975 |
| Map resolution (Å) | 3.30 | 3.75 | 3.07 | 3.16 | 2.83 | 4.52 |
| FSC threshold | 0.143 | 0.143 | 0.143 | 0.143 | 0.143 | 0.143 |
| **Refinement** |  |  |  |  |  |  |
| Initial model used (PDB code) | 7WLT | 7WLT | 7WLT | 7WLT | 7WLT | 7WLT |
| Model resolution (Å) | 3.3 | 3.8 | 3.0 | 3.2 | 2.8 | 4.6 |
| FSC threshold | 0.143 | 0.143 | 0.143 | 0.143 | 0.143 | 0.143 |
| Map sharpening *B* factor (Å<sup>2</sup>) | 79 | 96 | 52 | 52 | 63 | 108 |
| **Model composition** |  |  |  |  |  |  |
| Non-hydrogen atoms | 31695 | 31695 | 32145 | 32145 | 32145 | 32124 |
| Protein residues | 3852 | 3852 | 3915 | 3915 | 3915 | 3915 |
| Water | 0 | 0 | 0 | 0 | 0 | 0 |
| Ions | 0 | 0 | 0 | 0 | 0 | 0 |
| **B factors** |  |  |  |  |  |  |
| Protein | 182.42 | 192.52 | 144.47 | 142.11 | 143.26 | 717.62 |
| Ligand | / | / | / | / | / | / |
| Water | / | / | / | / | / | / |
| **R.m.s. deviations** |  |  |  |  |  |  |
| Bond lengths (Å) | 0.012 | 0.013 | 0.011 | 0.011 | 0.011 | 0.017 |
| Bond angles (°) | 1.172 | 1.184 | 1.139 | 1.137 | 1.130 | 2.002 |
| **Validation** |  |  |  |  |  |  |
| MolProbity score | 2.21 | 2.23 | 2.20 | 2.20 | 2.20 | 2.92 |
| Clash score | 15.69 | 16.01 | 15.98 | 15.38 | 15.92 | 59.00 |
| Poor rotamers (%) | 0.03 | 0.03 | 0.03 | 0.03 | 0.03 | 0.00 |
| **Ramachandran plot** |  |  |  |  |  |  |
| Favored (%) | 91.19 | 90.34 | 91.77 | 91.95 | 91.94 | 84.19 |
| Allowed (%) | 8.07 | 8.32 | 7.66 | 7.62 | 7.54 | 14.35 |
| Disallowed (%) | 0.75 | 1.34 | 0.58 | 0.53 | 0.52 | 1.47 |

## References

1 Levina, N. et al. Protection of Escherichia coli cells against extreme turgor by activation of MscS and MscL mechanosensitive channels: identification of genes required for MscS activity. EMBO J 18, 1730–1737, doi:10.1093/emboj/18.7.1730 (1999).

2 Sukharev, S. I., Blount, P., Martinac, B., Blattner, F. R. & Kung, C. A large-conductance mechanosensitive channel in E. coli encoded by mscL alone. Nature 368, 265–268, doi:10.1038/368265a0 (1994).

3 Coste, B. et al. Piezo1 and Piezo2 are essential components of distinct mechanically activated cation channels. Science 330, 55–60, doi:10.1126/science.1193270 (2010).

4 Maingret, F., Fosset, M., Lesage, F., Lazdunski, M. & Honore, E. TRAAK is a mammalian neuronal mechano-gated K+ channel. J Biol Chem 274, 1381–1387, doi:10.1074/jbc.274.3.1381 (1999).

5 Murthy, S. E. et al. OSCA/TMEM63 are an Evolutionarily Conserved Family of Mechanically Activated Ion Channels. Elife 7, doi:10.7554/eLife.41844 (2018).

6 Zhang, M. F. et al. Structure of the mechanosensitive OSCA channels. Nature Structural & Molecular Biology 25, 850-+, doi:10.1038/s41594-018-0117-6 (2018).

7 Zhang, M., Shan, Y., Cox, C. D. & Pei, D. A mechanical-coupling mechanism in OSCA/TMEM63 channel mechanosensitivity. Nat Commun 14, 3943, doi:10.1038/s41467-023-39688-8 (2023).

8 Bass, R. B., Strop, P., Barclay, M. & Rees, D. C. Crystal structure of MscS, a voltage- modulated and mechanosensitive channel. Science 298, 1582–1587, doi:DOI 10.1126/science.1077945 (2002).

9 Anishkin, A., Loukin, S. H., Teng, J. & Kung, C. Feeling the hidden mechanical forces in lipid bilayer is an original sense. Proc Natl Acad Sci U S A 111, 7898–7905, doi:10.1073/pnas.1313364111 (2014).

10 Martinac, B., Adler, J. & Kung, C. Mechanosensitive ion channels of E. coli activated by amphipaths. Nature 348, 261–263, doi:10.1038/348261a0 (1990).

11 Zhang, Y. et al. Visualization of the mechanosensitive ion channel MscS under membrane tension. Nature 590, 509–514, doi:10.1038/s41586-021-03196-w (2021).

12 Brohawn, S. G., Campbell, E. B. & MacKinnon, R. Physical mechanism for gating and mechanosensitivity of the human TRAAK K+ channel. Nature 516, 126–130, doi:10.1038/nature14013 (2014).

13 Schmidpeter, P. A. M. et al. Membrane phospholipids control gating of the mechanosensitive potassium leak channel TREK1. Nat Commun 14, 1077, doi:10.1038/s41467-023-36765-w (2023).

14 Lin, Y. C. et al. Force-induced conformational changes in PIEZO1. Nature 573, 230-+, doi:10.1038/s41586-019-1499-2 (2019).

15 Mulhall, E. M. et al. Direct observation of the conformational states of PIEZO1. Nature, doi:10.1038/s41586-023-06427-4 (2023).

16 Yang, X. et al. Structure deformation and curvature sensing of PIEZO1 in lipid membranes. Nature 604, 377–383, doi:10.1038/s41586-022-04574-8 (2022).

17 Zheng, W., Gracheva, E. O. & Bagriantsev, S. N. A hydrophobic gate in the inner pore helix is the major determinant of inactivation in mechanosensitive Piezo channels. Elife 8, doi:10.7554/eLife.44003 (2019).

18 Bae, C., Gnanasambandam, R., Nicolai, C., Sachs, F. & Gottlieb, P. A. Xerocytosis is caused by mutations that alter the kinetics of the mechanosensitive channel PIEZO1. Proc Natl Acad Sci U S A 110, E1162–1168, doi:10.1073/pnas.1219777110 (2013).

19 Zhou, Z. J. et al. MyoD-family inhibitor proteins act as auxiliary subunits of Piezo channels. Science 381, 799–804, doi:10.1126/science.adh8190 (2023).

20 Bae, C., Gnanasambandam, R., Nicolai, C., Sachs, F. & Gottlieb, P. A. Xerocytosis is caused by mutations that alter the kinetics of the mechanosensitive channel PIEZO1. P Natl Acad Sci USA 110, E1162–E1168, doi:10.1073/pnas.1219777110 (2013).

21 Glogowska, E. et al. Novel mechanisms of PIEZO1 dysfunction in hereditary xerocytosis. Blood 130, 1845–1856, doi:10.1182/blood-2017-05-786004 (2017).

22 Ma, S. et al. A role of PIEZO1 in iron metabolism in mice and humans. Cell 184, 969-+, doi:10.1016/j.cell.2021.01.024 (2021).

23 Ma, S. et al. Common PIEZO1 Allele in African Populations Causes RBC Dehydration and Attenuates Plasmodium Infection. Cell 173, 443–455 e412, doi:10.1016/j.cell.2018.02.047 (2018).

24 Goehring, A. et al. Screening and large-scale expression of membrane proteins in mammalian cells for structural studies. Nature Protocols 9, 2574–2585, doi:10.1038/nprot.2014.173 (2014).

25 Zhou, Z. et al. MyoD-family inhibitor proteins act as auxiliary subunits of Piezo channels. Science 381, 799–804, doi:10.1126/science.adh8190 (2023).

26 Song, Y. et al. The Mechanosensitive Ion Channel Piezo Inhibits Axon Regeneration. Neuron 102, 373–389 e376, doi:10.1016/j.neuron.2019.01.050 (2019).

27 Chambers, S. M. & Studer, L. Cell fate plug and play: direct reprogramming and induced pluripotency. Cell 145, 827–830, doi:10.1016/j.cell.2011.05.036 (2011).

28 Sukomon, N., Fan, C. & Nimigean, C. M. Ball-and-Chain Inactivation in Potassium Channels. Annual Review of Biophysics 52, 91–111, doi:10.1146/annurev-biophys-100322-072921 (2023).

29 Punjani, A., Rubinstein, J. L., Fleet, D. J. & Brubaker, M. A. cryoSPARC: algorithms for rapid unsupervised cryo-EM structure determination. Nat Methods 14, 290–296, doi:10.1038/nmeth.4169 (2017).

30 Pettersen, E. F. et al. UCSF Chimera--a visualization system for exploratory research and analysis. J Comput Chem 25, 1605–1612, doi:10.1002/jcc.20084 (2004).

